# Evolution of nasal and olfactory infection characteristics of SARS-CoV-2 variants

**DOI:** 10.1101/2022.04.12.487379

**Authors:** Mengfei Chen, Andrew Pekosz, Jason S. Villano, Wenjuan Shen, Ruifeng Zhou, Heather Kulaga, Zhexuan Li, Sarah E. Beck, Kenneth W. Witwer, Joseph L. Mankowski, Murugappan Ramanathan, Nicholas R. Rowan, Andrew P. Lane

**Author notes:** This work was funded by NIH Grants R01 AI132590, R01 DC016106 (A.P.L), by the Johns Hopkins Center of Excellence for Influenza Research and Surveillance, NIAID HHS N2772201400007C (AP), and the generosity of the collective community of donors to the Johns Hopkins University School of Medicine for COVID research. Corresponding Author: Andrew P. Lane, MD Department of Otolaryngology – Head and Neck Surgery Johns Hopkins Outpatient Center, 6^208^ floor, 601 N Caroline Street Baltimore, MD 21287-0910; Co-corresponding Author: Mengfei Chen, PhD, Department of Otolaryngology – Head and Neck Surgery Johns Hopkins University School of Medicine, 855 North Wolfe Street, Baltimore, MD 21205.

## Abstract

SARS-CoV-2 infection of the upper airway and the subsequent immune response are early, critical factors in COVID-19 pathogenesis. By studying infection of human biopsies in vitro and in a hamster model in vivo, we demonstrated a transition in tropism from olfactory to respiratory epithelium as the virus evolved. Analyzing each variants revealed that SARS-CoV-2 WA1 or Delta infects a proportion of olfactory neurons in addition to the primary target sustentacular cells. The Delta variant possesses broader cellular invasion capacity into the submucosa, while Omicron displays longer retention in the sinonasal epithelium. The olfactory neuronal infection by WA1 and the subsequent olfactory bulb transport via axon is more pronounced in younger hosts. In addition, the observed viral clearance delay and phagocytic dysfunction in aged olfactory mucosa is accompanied by a decline of phagocytosis related genes. Furthermore, robust basal stem cell activation contributes to neuroepithelial regeneration and restores ACE2 expression post-infection. Together, our study characterized the nasal tropism of SARS-CoV-2 strains, immune clearance, and regeneration post infection. The shifting characteristics of viral infection at the airway portal provides insight into the variability of COVID-19 clinical features and may suggest differing strategies for early local intervention.

## Introduction

Severe acute respiratory syndrome coronavirus 2 (SARS-CoV-2), the causative pathogen in the worldwide pandemic of coronavirus disease 2019 (COVID-19), is readily transmitted via respiratory droplets during close contact. The nasal cavity is the entry point of respiratory tract, and the high viral load detected there indicates that this is the principal initial site of SARS-CoV-2 infection and immune response^1, 2^. The loss of the sense of smell is common in patients infected with the SARS-CoV-2 original WA1 and later Delta strains. Pathological studies visualizing olfactory viral infection in postmortem samples of nasal respiratory and olfactory epithelium partially explain the smell loss in COVID-19 patients^3^. Unlike the previous strains, Omicron rarely causes olfactory loss, possible suggesting a change in cellular tropism as the virus evolved. Systematic characterization of the SARS-CoV-2 infection pattern in the nose is important for understanding COVID-19 pathogenesis and for developing early local intervention.

The cellular tropism of SARS-Cov-2 in the nasal cavity is relevant to pathologic tissue damage and to COVID testing. Cellular entry of SARS-CoV-2 depends on the binding of the virus spike S protein to angiotensin-converting enzyme 2 (ACE2) in host tissue^4, 5^. The level of viral receptors and its subcellular localization is a key determinant of susceptibility to infection. In parallel with the gradually decreased ACE2 RNA expression pattern from the upper airway to distal intrapulmonary regions^6, 7^, *in vitro* SARS-CoV-2 infection of human respiratory epithelial cell cultures shows a gradient of diminishing infectivity (olfactory epithelial cells were not included in these studies)^7^. Notably, we have observed up to 700-fold higher expression of ACE2 in the sustentacular cells of olfactory epithelium in comparison to respiratory epithelial cells in human nose and trachea^8^. Whether the enrichment of ACE2 in the olfactory epithelium correlates with more susceptibility to SARS-CoV-2 infection than respiratory cells, and how the infection affects the olfactory sensory neurons are largely unknown.

Despite the substantially reduced COVID-19 incidence in adults after the rapid vaccination program rollout, the unvaccinated population including young children, as well as break-through infections by new variants, now comprise the majority cases. SARS-CoV-2 infection typically causes mild acute airway illness in approximately 81% of COVID-19 patients; however, 14-17% of hospitalized cases experience severe symptoms and require intensive care^9^. The severity of COVID-19 is highly age-related, with a fatality rate that can reach 30.5% in patients of 85 years or older^10^, suggesting compromised anti-viral immunity with aging. The correlation between aging and the cellular damage and subsequent immune response to SARS-CoV-2 requires further investigation.

In this study, we perform *in vitro* infection of human nasal explants and show an extremely high infection rate of SARS-CoV-2 WA1 strain in olfactory epithelium relative to the adjacent respiratory epithelium. By comparing the infection patterns of WA1, Delta, and Omicron strains in the hamster nasal cavity, we demonstrated a transition in tropism from olfactory to respiratory epithelium as the virus evolved, providing insight into COVID-19 pathogenesis and diagnosis. Using a WA1 strain infected hamster model, our additional findings demonstrate an age-associated infection of olfactory neurons and impaired macrophage phagocytosis. These findings indicate that the nasal viral replication and local immune defense could be a potential target of early intervention.

## Results

### WA1 strain primarily targets human olfactory neuroepithelium

The human olfactory mucosa is located in the superior part of the nasal cavity contains sustentacular cells and olfactory sensory neurons that is responsible for the sense of smell. To examine the precise cellular tropism of SARS-CoV-2 in the nasal cavity including both respiratory and olfactory epithelium, we initially performed WA1 strain *in vitro* infection experiments using human nasal tissue discarded in endonasal sinus and skull base surgery in COVID-19-negative individuals. To establish a reliable protocol to detect SARS-CoV-2 antigen, we screened and verified 4 different antibodies for visualizing spike (S) or nucleocapsid protein (NP) in the infected tissue sections (Extended Data Fig. 1a). The staining pattern of antibodies predominant located in apical sustentacular cells is consistent with the viral RNA detected by RNAscope analysis (Extended Data Fig.1b).

**Fig. 1.**
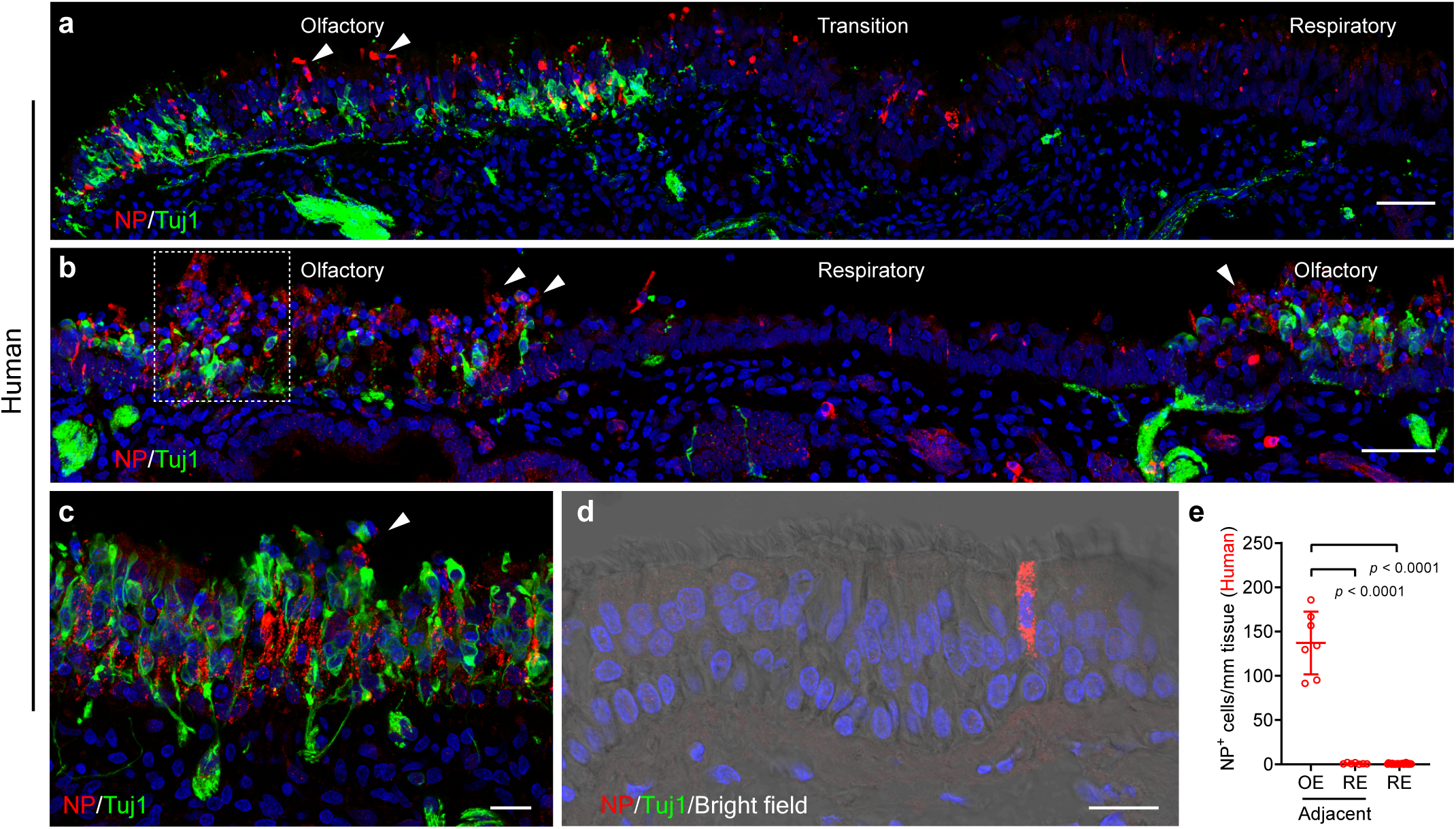
SARS-Cov-2 WA1 selectively targets human olfactory neuroepithelium. **a**,**b**, Confocal images of SARS-CoV-2 viral antigen NP (red, Novus, NB100-56576) and olfactory neuronal marker β-III Tubulin (Tuj1, green) in superior turbinate biopsies from 2 separate patients. Images were obtained under tile scan mode, which covered olfactory and adjacent respiratory epithelium in the same piece of tissue. Boxed area in (**b**) was highlighted in Extended Data Fig. 1c, d. **c**, Co-staining of NP and Tuj1 in human biopsy collected from the olfactory cleft. **d**, Representative image of NP overlapped with Tuj1-negative ciliated cell (brightfield). Confocal image was obtained from a biopsy which contains only respiratory epithelium. **e**, Quantification of NP^+^ cells per mm tissue. 24 independent specimens have exclusively respiratory epithelium (RE), while 7 specimens contained both respiratory and olfactory epithelium (OE). Arrowheads (**a**-**c**) indicate the detachment of infected cells. Data in (**e**) are represented as mean ± S.D. *p* value was calculated by one-way ANOVA. Scale bars, 50 µm (**a** and **b**); 20 µm (**c,d**).

Because the olfactory mucosa is irregularly distributed and surrounded by respiratory epithelium in the human nasal cavity ^11^, we included the neuronal marker Tuj1 for the verification of olfactory epithelium. By immunostaining with SARS-CoV-2 NP, we observed substantial viral antigen in the Tuj1^+^ olfactory region at 9 hours post infection, but very little NP in the adjacent Tuj1 negative respiratory epithelium (Fig. 1a,b). The vast majority of NP^+^ cells co-localized with Krt18^+^ olfactory sustentacular cells (Extended Data Fig. 1c,d). Viral infection caused extensive sustentacular cell death, with rapid detachment and sloughing into the nasal lumen (Fig. 1a,b, and Extended Data Fig. 1c,d). Compared to the mock control (Extended Data Fig. 1e), structural damage was readily detected in the olfactory mucosa but not in the respiratory epithelium (Fig. 1a, b). A high viral infection ratio was also found in human olfactory cleft specimens obtained from skull base surgery (Fig. 1c). Low viral infection was observed in 24 explants that only contained respiratory epithelium (Fig. 1d,e). We quantified the number of NP^+^ cells in 7 tissue explants, revealing 100-300-fold more infected cells in olfactory epithelium compared to adjacent respiratory epithelium (Fig. 1e). These results together our earlier observed enrichment of ACE2 expression illustrated an olfactory specific tropism of SARS-CoV-2 WA1 strain and explained the common symptom of anosmia in COVID-19 patients.

### Omicron variant shows transition in tropism from olfactory to respiratory epithelium

As SARS-CoV-2 evolved, new variants including Delta and Omicron caused surges in cases worldwide. The tropism of these different strains in the nasal cavity has not been clarified. We next characterized the cellular tropism of WA1, Delta, and Omicron in nasal mucosa using a hamster model. These experiments allowed us to determine whether the observed cellular tropism of WA1 in human olfactory epithelium is applicable to new variants in animal models and relates to subsequent disease pathogenesis.

The olfactory mucosa in the hamster nasal cavity is located in the posterior and dorsal aspect, while the anterior and ventral areas are respiratory. We confirmed that the expression of the neuronal marker Tuj1 in olfactory epithelium was mutually exclusive with the respiratory marker Foxj1 in the nasal cavity (Extended Data Fig. 2a). Therefore, the olfactory epithelium can be identified based on Tuj1 positive staining, the presence of axon bundles, and the relatively increased thickness of the neuroepithelium. After SARS-CoV-2 inoculation (1×10^5^ TCID50), we captured confocal images of the entire nasal cavity in coronal sections at three different levels (Fig. 2a). At 4 days post infection (dpi), we verified the extremely high viral antigen NP in the Tuj1^+^ olfactory epithelium of WA1 or Delta-infected hamsters, with a sharp decline in the adjacent respiratory epithelium (Fig. 2b). About 79.2% or 70.3% length of Tuj1^+^ olfactory epithelium was infected by WA1 or Delta, respectively (Fig. 2b, c). We observed that the expression of ACE2 in some OMP^+^ olfactory areas is low or undetectable, interpretating the uninfected areas in WA1 or Delta treated groups (Extended Data Fig. 2b).

**Fig. 2.**
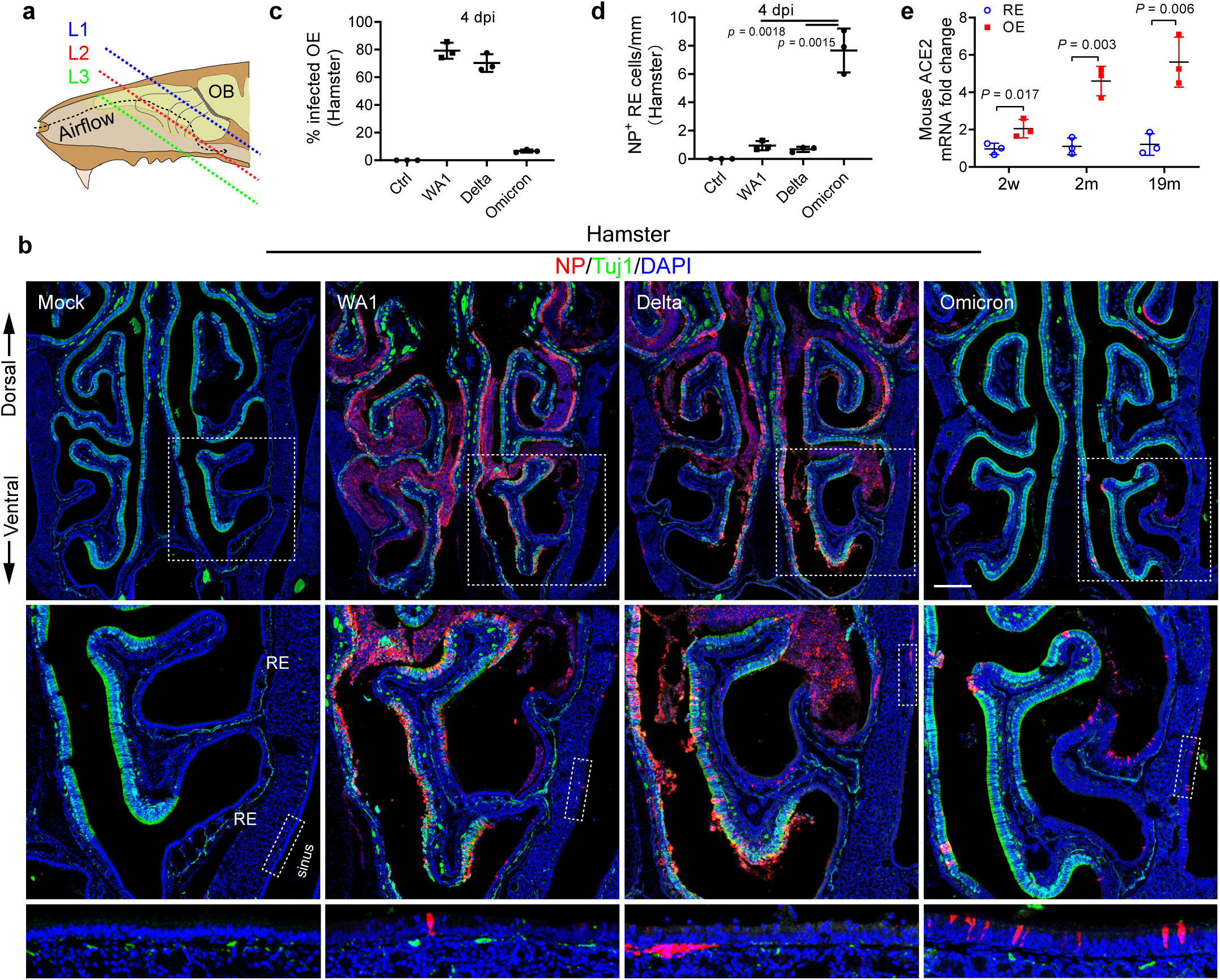
Omicron variant shows tropism transition from olfactory to respiratory epithelium. **a**, Scheme of the tissue section. To avoid variability across different animals, frozen sections were collected and examined at three consistent levels (L1–3) representing the anterior (mainly respiratory epithelium), middle (respiratory + Olfactory epithelium), and posterior (mainly olfactory epithelium). **b**, Confocal images of NP and Tuj1-labeled hamster nasal sections at L2. WA1, Delta, and Omicron infected hamsters were examined on 4 dpi. Boxed areas are highlighted at bottom. Scale bars = 500 µm. **c**, Percentage of the infected olfactory epithelium. The total length of Tuj1^+^ or NP^+^/Tuj1^+^ epithelium in each section at L1-3 were quantified using Image J. **d**, Quantification of NP^+^ cells in nasal respiratory epithelium. The total NP^+^ cells in Tuj1^-^ respiratory epithelium including paranasal sinuses of each section were counted. **e**, qPCR analysis of ACE2 expression in mouse nasal respiratory or olfactory epithelium at age of 2 weeks, 2months, and 19months. The entire nasal respiratory or olfactory epithelium from the same animal were isolated separately. Data are represented as mean ± S.D. Statistical significance was determined by unpaired two-tailed *t*-test. Each data point represents an individual animal.

In contrast to WA1 or Delta strain, the infected olfactory epithelium in Omicron group was dramatically reduced to 6.7%, which is consistent with earlier reports of a comparatively lower pathogenicity in lung of Omicron infected hamsters^12, 13^. The low infection rate of Omicron in olfactory epithelium (Fig. 2b, c) seems to correlate with the low incidence of smell loss in patients. Interestingly, we observed the Omicron infected NP^+^ nasal and sinus respiratory cells was increased 7-10-fold when compared to WA1 or Delta, suggesting an olfactory to respiratory tropism transition with the Omicron variant (Fig. 2b, d). These tropism patterns were further demonstrated in sections of the anterior or posterior nasal cavity where the proportion of respiratory epithelium is much greater or less, respectively (Extended Data Fig. 2c, 3a). Together, these results identify that the SARS-Cov-2 variants have different tropism in nasal mucosa that may play a role in the shifting pathogenic features of COVID-19 as the virus evolved.

**Fig. 3.**
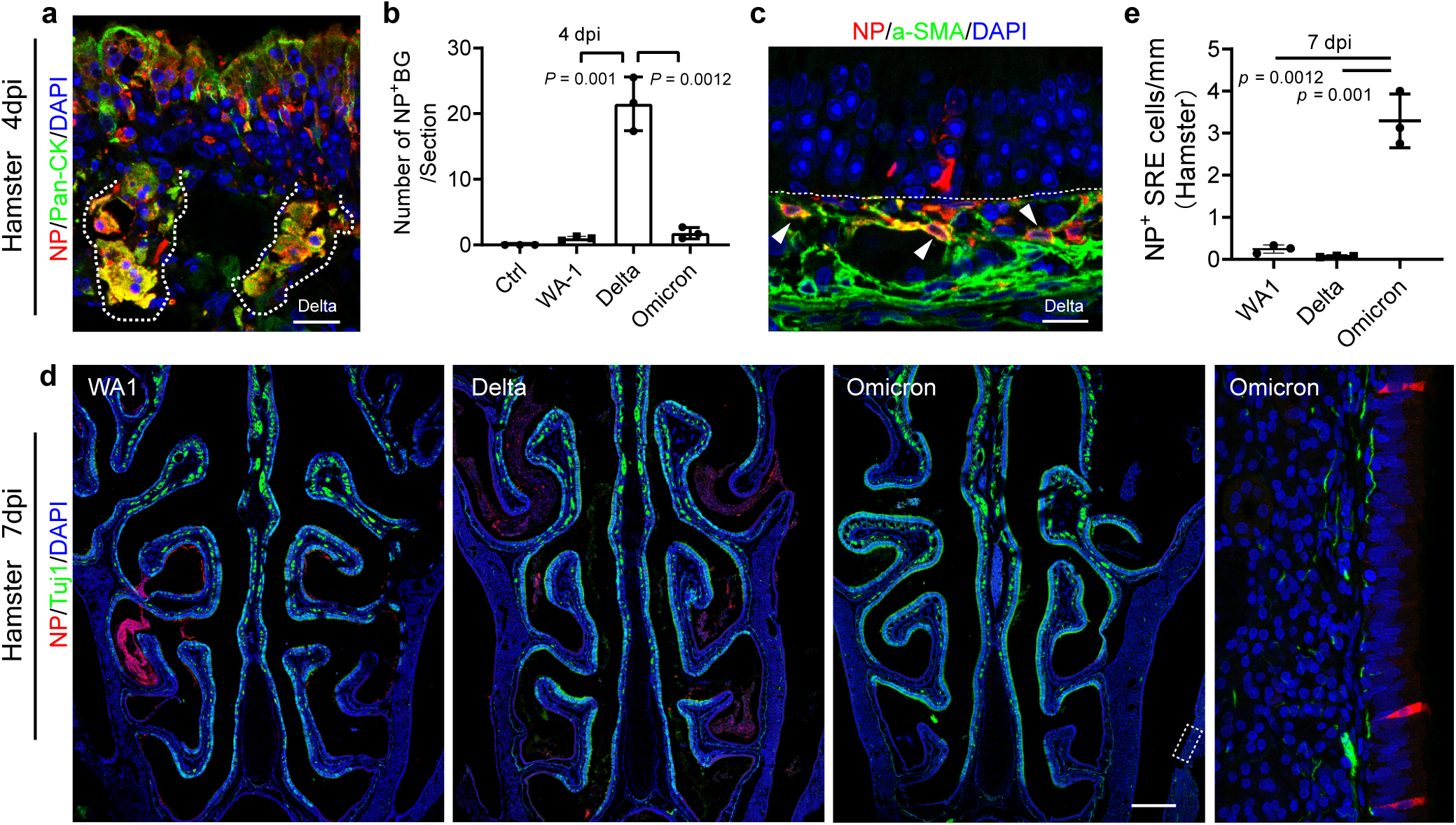
Delta variant infects cells in submucosa of the nose. **a**, Representative image shows NP^+^/Pan-Keratin^+^ Bowman’s glands in Delta treated hamsters. **b**, Quantification of infected Bowman’s glands. The average number of NP^+^ Bowman’s glands in one 14 µm section was calculated. 3 sections per animal were counted. **c**, Confocal image shows NP^+^/ 4dpi were examined. α-SMA^+^ myofibroblasts. Hamsters infected with Delta variant on 4dpi were examined. **d**, Co-staining of Tuj1 and NP in nasal sections at 7dpi. Whole nasal cavity images were captured using a tile scan and z stack mode on a 14 µm section. Boxed area in Omicron infected hamster is highlighted on the right. Scale bars = 500 µm. **e**, Quantification of NP^+^ respiratory epithelial cells in paranasal sinuses. 3 sections per animal was counted. Data are represented as mean ± S.D. Statistical significance was determined by unpaired two-tailed *t*-test. Each data point represents an individual animal.

To support the olfactory epithelial tropism of WA1 and Delta, we further performed qPCR analysis of ACE2 expression in the entire nasal respiratory or olfactory mucosa in C57BL/6J wildtype mouse at the ages of 2 weeks, 2 months, and 19 months. Compared to the nasal respiratory epithelium, ACE2 mRNA transcription in adult olfactory epithelium was increased 5-7-fold in 2m or 19m old animals (Fig. 2e). It should be noted that ACE2 mRNA levels in whole olfactory mucosa are greatly diluted by the larger proportion of ACE2-low-to-negative cells (neurons), relative to respiratory mucosa. In addition, ACE2 protein in human^14, 15^ or mouse^16^ epithelial tissue is predominantly expressed at the apical surface. The more diffuse cellular pattern of ACE2 staining in human autopsy specimens may result from post-mortem degradation.

In any putative human olfactory tissue sample, a neuronal marker must be utilized for verification because the olfactory mucosa is irregularly distributed and surrounded by respiratory epithelium. Consistent with previous data, we observed a gradually increased ACE2 expression in olfactory mucosa from 2 weeks through adulthood^17, 18^. The level of ACE2 expression in nasal respiratory mucosa was comparable between young and old animals (Fig. 2e). Together, the ACE2 expression pattern supports the olfactory epithelium as a site of SARS-CoV-2 replication especially for WA1 or Delta variants. The decreased olfactory tropism in the Omicron variant is consistent with the recently reported endocytic entry pathway^19, 20^.

### Delta variant demonstrates greater infection of cells in the nasal submucosa

In the lamina propria, we frequently detected NP^+^ cells in Delta inoculated hamsters at 4dpi. Co-staining of NP and Pan-cytokeratin revealed that some of those infected cells were Bowman’s glands (Fig. 3a), the producer of specialized mucus critical for odor perception^21^. These results are in line with our previously reported ACE2 expression in human biopsies^8^ and the observation that SARS-CoV-2 targets Bowman’s glands in postmortem samples by other groups^3^. The number of NP^+^ Bowman’s glands in Delta infected hamsters increased 21-fold when compared to WA1, and is sharply decreased in Omicron group (Fig. 2b, 3b). Additionally, NP^+^ elongated submucosal cells can be readily detected in olfactory and respiratory mucosa of Delta-infected animals (Fig. 3c) but is dramatically reduced in Omicron treated hamsters. These NP^+^ cells are aSMA^+^ but negative for Iba1 (macrophage marker) and Vimentin (mesenchymal and olfactory ensheathing cell marker), suggesting the contractile myofibroblasts/mesenchymal cell lineage (Fig. 3c, Extended Data Fig. 3b). The broader cell types targeted in the submucosa by the Delta variant may increase the severity of tissue damage.

We next asked whether the infected submucosal cells are rapidly cleared or instead serve as an ongoing viral reservoir. At 7dpi, we observed almost all the NP^+^ olfactory epithelial cells had been lost, other than those in sloughed off debris in the nasal lumen. In the submucosa, except NP^+^ axon in WA1 group, NP^+^ cell was barely detectable in animals infected with any of the 3 strains (Fig. 3d). These results in agreement with the reported viral titer analysis at 7dpi^12, 13^. However, in the paranasal sinuses, an area was not examined in earlier studies^12, 13^, we detected a small number of NP^+^ respiratory epithelial cells in WA1 but rarely in Delta treated hamsters at 7dpi. In parallel with the tropism transition from olfactory to respiratory epithelium, more pronounced NP^+^ sinonasal epithelial cells (3.3 positive cells/mm epithelium) were observed in Omicron variant-treated hamsters (Fig. 3d, e), suggesting a longer duration of the Omicron variant infection in sinus epithelium relative to the ancestral SARS-CoV-2 strains. It is unknown whether those Omicron -infected cells in the sinuses are actively transmitting virus at 7dpi.

### Age associated SARS-CoV-2 WA1 infection of olfactory sensory neurons

While neurological symptoms, including headache, encephalitis, and altered mental status have been reported in COVID-19 patients ^22, 23^, the evidence of SARS-CoV-2 olfactory neuronal infection is controversial^3, 24^. Earlier studies have shown SARS-CoV-2 RNA or viral antigen in postmortem brain tissue samples^25, 26^, and rare infection observed in olfactory neurons in autopsy tissue hints towards transmucosal invasion^27^. The reported data have indicated SARS-CoV-2 infection affects neurons in the hamster model^28, 29^; however, Tuj1^+^ immature neurons are normally located next to the basal layer, and the long foot-like processes of infected sustentacular cells surrounding olfactory neurons could be mis-interpretated in earlier reports. The direct evidence of olfactory neuronal infection and the factors that affect the frequency of infection and entry to the brain remain to be clarified^3, 24^.

Given the high tropism of SARS-CoV-2 WA1 or Delta in olfactory mucosa, we took advantage of a hamster model to examine WA1 or Delta infection in the olfactory neuronal population. The hamster model allowed us to avoid the significant limitations of autopsy tissue, including an often prolonged and severe disease course and tissue degradation during the postmortem interval. We utilized a higher viral inoculum (1×10^7^ TCID50) to generate more uniform infections that would allow us to identify variation across age groups^30^. As expected, we observed the vast majority of NP^+^ cells were apical sustentacular cells^31^ in WA1 infected hamsters (Extended Data Fig.1a,b) at 4 dpi. Interestingly, in the superior turbinate of posterior nasal cavity, we observed NP labeling of a small portion of cells located in the olfactory sensory neuronal layer and their axon bundles (Fig. 4a). Co-staining of NP with neuronal markers Tuj1(immature) and OMP (mature) revealed viral infection in a subset of cells from the neuronal lineage (Fig. 4b, c). NP^+^/OMP^+^ infected olfactory neurons were also detected in Delta variant treated hamsters (Extended Data Fig. 4a). We detected viral antigen travel along the Tuj1^+^ axon from epithelium to the lamina propria (Fig. 4d, e). In axon bundles, NP co-localized with Tuj1^+^ or OMP^+^ axons (Extended Data Fig. 4b, c) but did not colocalize with Vimentin^+^ ensheathing cells (Extended Data Fig. 4d). In addition, we confirmed the olfactory neuronal infection by WA1 or Delta at 1×10^5^ TCID50 (Extended Data Fig. 4e, f). Precise quantification of the number of infected olfactory neurons is a challenge because the intensity of marker staining in infected and dying cells subsides^3^ compared to normal cells (Extended Data Fig. 4g) and because the epithelium sloughs off in some areas. We observed at least 20 NP^+^/OMP^+^ or Tuj1^+^ neurons in each section of hamster infected with the WA1 at 1×10^5^ TCID50. Compared to WA1, olfactory neuronal infection is sharply decreased in Delta and rare in Omicron group. These data suggested that WA1 or Delta can also infect a proportion of olfactory sensory neurons, in addition to sustentacular cells that are the primary target in the upper airway. We therefore used WA1 strain for the following aging-related experiments.

**Fig. 4.**
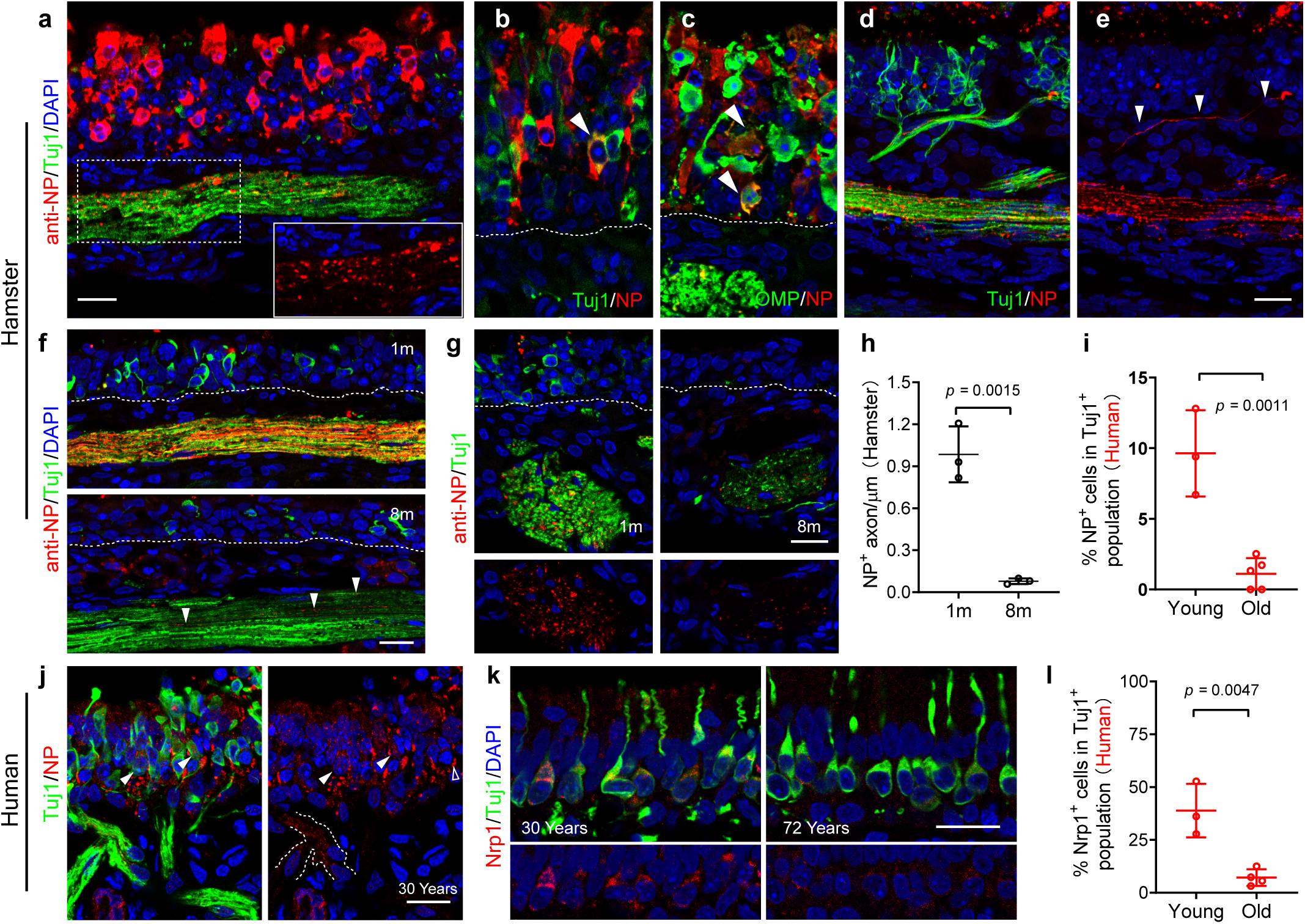
Age associated SARS-Cov-2 infection in olfactory sensory neurons. **a**-**c**, Confocal image showing WA1-infected hamster olfactory epithelium at 4dpi. Insert in (**a**) highlighting an NP stained axon bundle (horizontal section). Arrowheads indicate virus infected Tuj1^+^ immature (**b**) or OMP^+^ mature (**c**) sensory neurons (coronal sections). White line indicated the basal layer of epithelium. **d**, **e**, NP^+^ axon travel from neuroepithelium to laminar propria and merge into Tuj1^+^ axon bundle. **f-h**, Quantification of NP^+^ axons in young and old hamsters at 6dpi. Representative images show horizontal (**f**) or coronal sections (**g**). NP^+^ axons were quantified per µm of the diameter of axon bundle. **i**,**j**, Representative images showing NP located in Tuj1^+^ human olfactory neurons (**j**) and the percentage of NP^+^ cells in Tuj1^+^ population (**i**). Dotted line in (**j**) indicates virus infected NP^+^ axon. Arrowheads denote NP^+^/Tuj1^+^ neurons compared to uninfected cells (empty arrowhead). Infected biopsies from 3 young donors (age 25-33 years) and 5 biopsies from older donors (age 54-72 years) were quantified for Tuj1^+^ neuronal infection. **k**,**l**, Representative images of Nrp1 expression in human olfactory epithelium (**k**) and quantification of Nrp1^+^ cells in Tuj1^+^ population (**l**). 3 biopsies from young (age 20-30 years) and 4 biopsies from older donors (age 68-79 years) were examined for Nrp1 expression. Images in (**f**) were captured with 3 µm Z-stack and exported by maximum intensity projections. Each data point represents an individual sample from hamster (**h**), or human (**i** and **l).** Details of human biopsies can be found in Supplementary Table 1. Data are represented as mean ± S.D. Statistical significance was determined by unpaired two-tailed t-test. Scale bars, 20 µm.

The rare expression of ACE2 in olfactory sensory neurons^8, 16^ suggests that neuronal entry may mediated by other receptors such as Neuropilin-1(Nrp1)^32, 33^. In the olfactory epithelium, Nrp1 was expressed in the olfactory nerve in the embryonic stage and in immature neurons after birth^34, 35^. By using qPCR analysis, we detected 2.7-fold reduction of Nrp1 mRNA in the olfactory epithelium of 19-month-old compared to 2 weeks young mice (Extended Data Fig. 5a). Age related Nrp1 reduction in the olfactory epithelium was also verified by immunohistochemistry. About 34.2% of Tuj1^+^ olfactory neurons express Nrp1 in young mice but only 9.7% of Tuj1^+^ neurons in the aged group display a low level of Nrp1(Extended Data Fig. 5b-d). A few mature olfactory neurons in young mice also express Nrp1 (Extended Data Fig. 5b). In addition, Nrp1 can be detected in the axon bundles and periglomerular cells in young olfactory bulb, but are sharply declined in aged mice (Extended Data Fig. 5b,e).

**Fig. 5.**
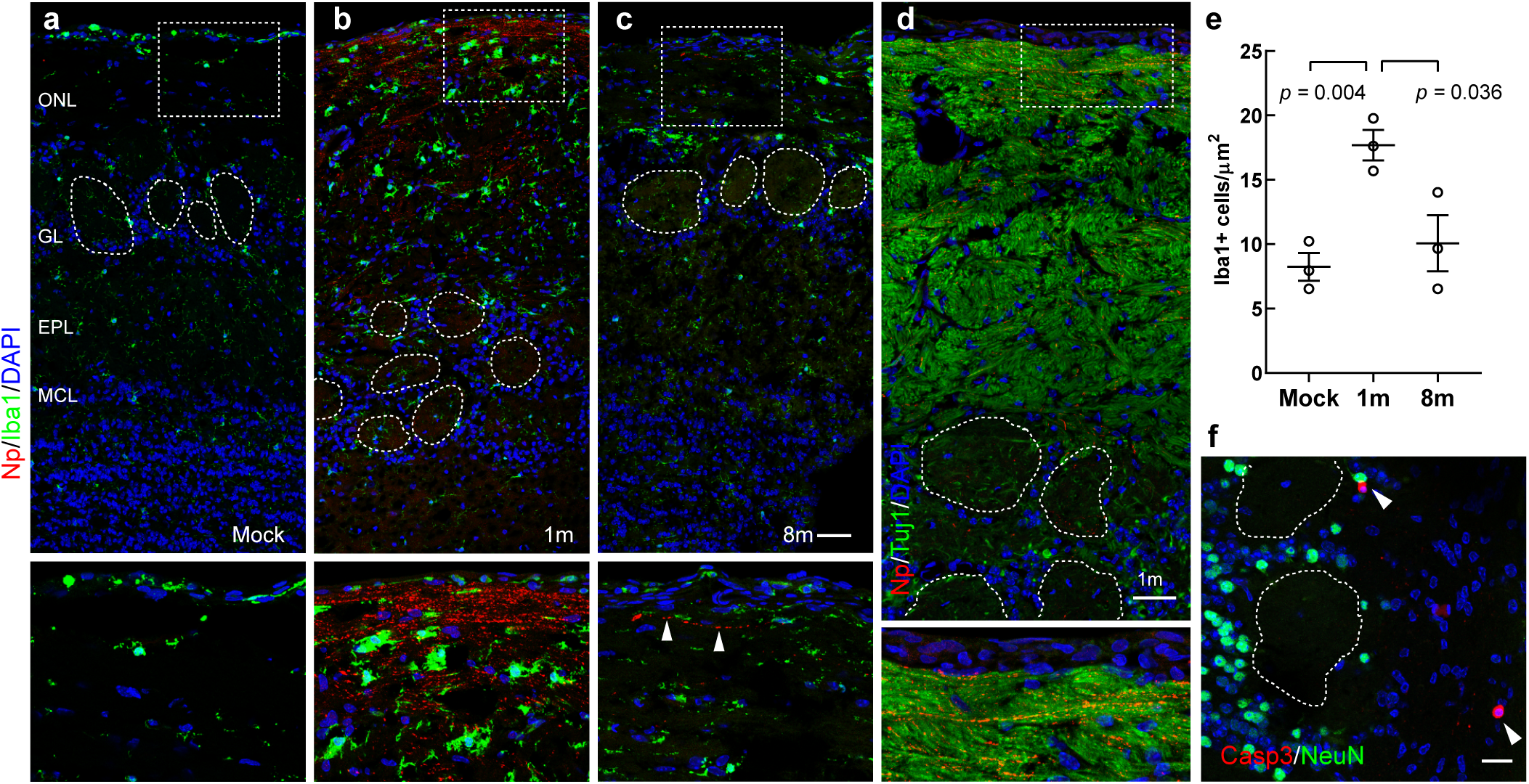
Increased olfactory bulb transport of SARS-CoV-2 in young hamsters. **a**-**c**, Confocal images of Iba1 and NP co-staining in hamster olfactory bulbs. Arrowheads indicate infected axon. **d,** Co-staining of NP and Tuj1 in a serial section next to panel (**b**). **e**, Quantification of Iba1^+^microglials in hamster olfactory bulb. Each data point in (**d**) represents an individual hamster sample. Data are represented as mean ± S.D. Statistical significance was determined by unpaired two-tailed t-test. **f**, Confocal image of cleaved caspase-3^+^/NeuN^-^ apoptotic cells (arrowheads) in the glomerular layer at 4dpi. Images were captured with 3 µm (**a**-**d**) or 4 µm (**f**) Z-stack and exported by maximum intensity projections. Olfactory bulb tissues were collected from young and old hamsters on 6dpi (**a**-**d**) or from mock control. Scale bars, 50 µm, (**a**-**d)**; 20 µm, (**f**). ONL, olfactory nerve layer; GL, glomerular layer; EPL, external plexiform layer; MCL, mitral cell layer. Boxed areas are highlighted at bottom. Dotted circles indicate glomeruli.

The age-related pattern of Nrp1 expression indicated a potential higher efficiency of SARS-CoV-2 infection in olfactory neurons in young population. To assess whether age could be a factor mediating neuronal infection in the olfactory epithelium, we performed SARS-CoV-2 WA1 (1×10^7^ TCID50) infection experiments using young (1-month) and aged (8-month) hamsters. At 6 dpi, viral antigen (NP) could be readily detected in the axon bundles in young hamsters, but infected axons were dramatically decreased in older hamsters (Fig. 4 f-h). We also examined the WA1-infected human explants and identified a remarkable increase of viral load in Tuj1^+^ neurons and axon bundles in tissue from young individuals (<30 years old) (Fig. 4i, j). As expected, we observed 38.9% of Tuj1^+^ olfactory neurons co-express Nrp1 in younger human biopsies, but the proportion of Tuj1^+^/Nrp1^+^ neurons dramatically reduced (7.2%) in older adults (Fig. 4k, l). Together, these results support age-dependent olfactory neuron infection and axonal transport.

### Increased olfactory bulb axonal transport of WA1 in young hamsters

The increased frequency of viral NP in the axons of younger animals indicated that SARS-CoV-2 WA1 may be prone to accessing the brain in this population. To verify this hypothesis, we examined the olfactory bulbs of 1 and 8-month old hamsters. At 6 dpi. we detected NP^+^ axons located in the olfactory nerve layer (ONL) in young hamsters (Fig. 5a,b), suggesting the viral transport to olfactory bulb. Compared to the young hamsters, infected axons are rarely detected in the older group (Fig. 5a-c). Co-staining analysis of serial sections verified that the NP signal is located in the Tuj1^+^ olfactory nerve layer (Fig. 5d). In the leptomeningeal layer where the viral RNA signal was detected in postmortem samples^3^, the NP antibody staining was not detectable in hamster (Fig. 5a-d). In addition, the observed leptomeningeal viral RNA staining was speculated to be extracellular virions instead of intracellular viral RNA synthesized by infected cells^3^. In parallel to the greater olfactory bulb viral transport, the number of Iba1^+^ microglia cells in young olfactory bulb was increased 1.7-fold compared with older group (Fig. 5e). No viral antigen could be detected in the mock control.

Immunostaining of horizontal sections crossing the olfactory mucosa and forebrain region revealed a massive number of NP^+^ axons traveling from the lateral olfactory epithelium to olfactory bulb in young, but not aged, hamsters at 6 dpi (Extended Data Fig. 6a-f). In line with the reported Nrp1 expression in lateral olfactory nerve, which contains axons from turbinate neurons^36^, the infected axon in the septum nerve was rare. NP^+^ axons could also be detected in glomeruli where the olfactory sensory neuron axon terminal projections synapse with OB mitral cells (Fig. 5b, d) at 6dpi. As a consequence of olfactory viral transport, we observed Caspase-3^+^ apoptotic cells and virus RNA in the glomerular layer at 4dpi (Fig. 5f, and Extended Data Fig. 6g,h) in the young group. These Caspase-3^+^ cells were negative for Iba1 or the neuronal marker NeuN. The transported virus in olfactory bulb appears to lose the capacity for replication based on the restriction of NP signal to axons in the outer olfactory nerve layer and glomeruli at 6dpi (Fig. 5b-d). Despite the close anatomic relationship between the olfactory mucosa and the nearby OB axons, no obvious transmucosal viral antigen NP was displayed except within axons.

**Fig. 6.**
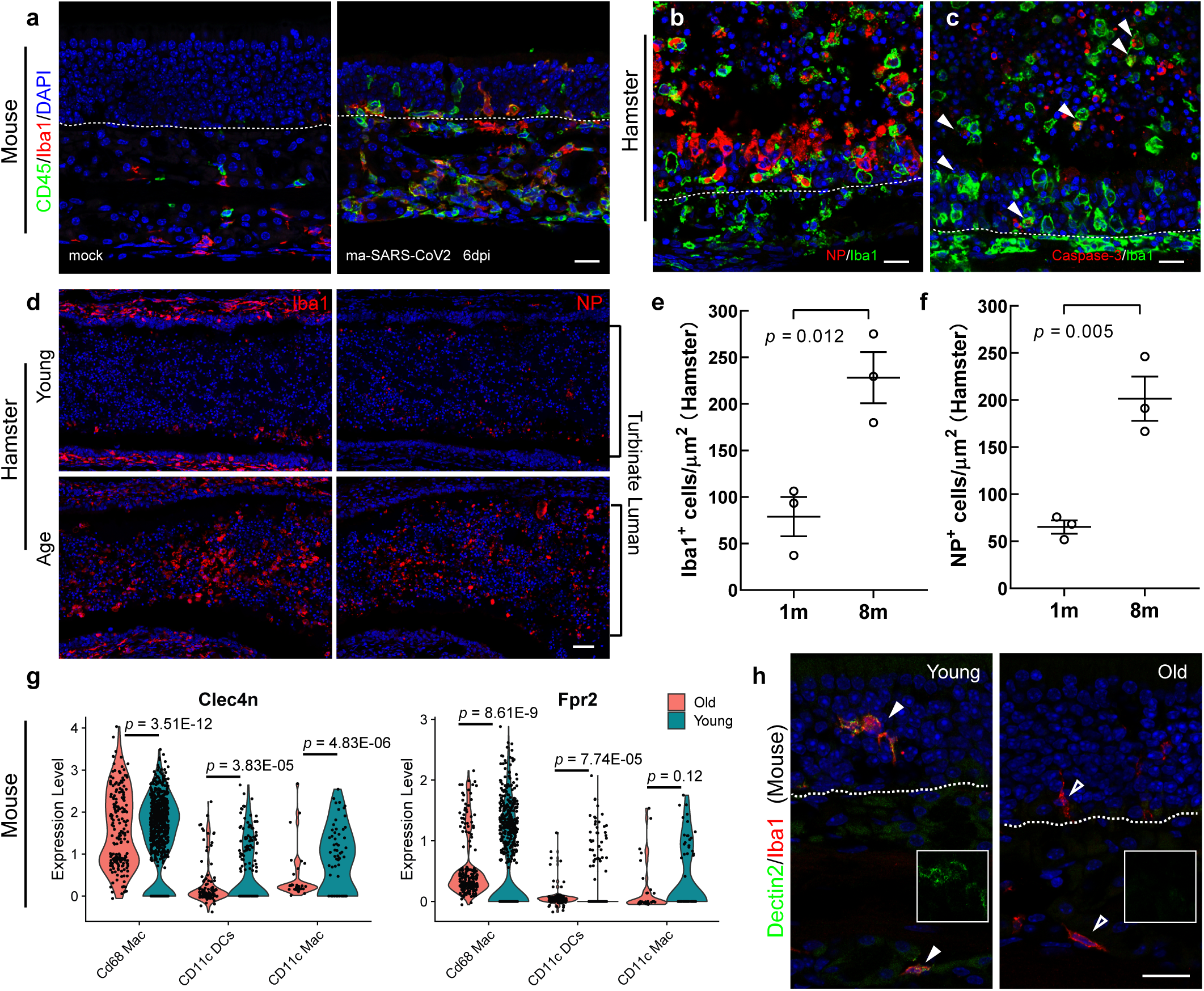
Age-associated delay in viral clearance in olfactory mucosa. **a**, Representative images showing CD45 and Iba1 co-staining in olfactory mucosa. Mock or maSARS-Cov2 infected wildtype mice were examined at 6dpi. **b**, Co-immunostaining shows Iba1^+^ macrophages engulfing NP^+^ debris in hamster olfactory mucosa at 4dpi. **c**, Representative image of Iba1 and cleaved caspase-3 co-staining in hamster at 4dpi. Arrowheads highlight the Iba1^+^ macrophages undergoing apoptosis. **d**, Representative images showing Iba1 or NP staining in serial sections. Each panel combines 6 40x images acquired under tile scan mode. Young or old hamsters’ olfactory tissues were examined at 6dpi. **e**,**f**, Quantification of Iba1^+^ (**e**) or NP^+^ (**f**) cells in hamster nasal olfactory lumen at 6dpi. Serial sections (**d**) from 4 different levels were quantified. **g**, Violin plots showing the differentially expressed Clec4n (Dectin2) or Fpr2 in young and old macrophage/dendritic lineage. **h**, Confocal images of Iba1 and Dectin2 co-staining in mouse olfactory mucosa. Each data point represents an individual hamster sample. Statistical significance was determined by unpaired two-tailed t-test. The white dotted line in (**a**-**c**) indicates the basement membrane. Scale bars, 20 µm (**a**-**c**, **h**); 50 µm (**d**).

Similar to ACE2 expression in lung vascular endothelial cells^37^, ACE2 in the mouse or hamster olfactory bulb is mainly located in the blood vessels (Extended Data Fig. 6i,j). We observed CD45^+^/Iba1^-^ immune cells infiltrating into the olfactory bulb in SARS-Cov2 infected hACE2 mice (Extended Data Fig. 6k,l), indicating passage of leukocytes across an impaired blood-brain barrier. Given the lack of lymphatic vessels in brain parenchyma, it is unlikely that viral infection of the olfactory bulb occurs via this route^38^. The inflammatory response in the hamster brain is not as severe as in the hACE2 mouse model, therefore the vascular damage is also likely much milder in hamster. Together, these results support that SARS-CoV-2 WA1 can gain access to the olfactory bulb region in the brain mainly through olfactory neuronal axons with higher frequency in younger population, while virus replication is limited.

### Age-related viral clearance delay and phagocytic dysfunction in the olfactory mucosa

The tropism of SARS-CoV-2 in olfactory epithelium indicates the capacity of local immune system against viral infection could involve in the pathogenesis of COVID-19. It has been reported that reduced innate antiviral defenses including type I and type III interferons coupled with a hyperinflammatory response is the major cause of disease severity and death in COVID-19 patients^39, 40^. Corresponding to the high viral load in olfactory epithelium, our qPCR analysis revealed an extensive upregulation of the anti-viral gene Ifng (type II interferon) in the nasal turbinate tissue post infection (Extended Data Fig. 7a), suggesting activated local immune defense. We next studied the potential age-related alternation of olfactory immune response to SARS-CoV-2 infection.

**Fig. 7.**
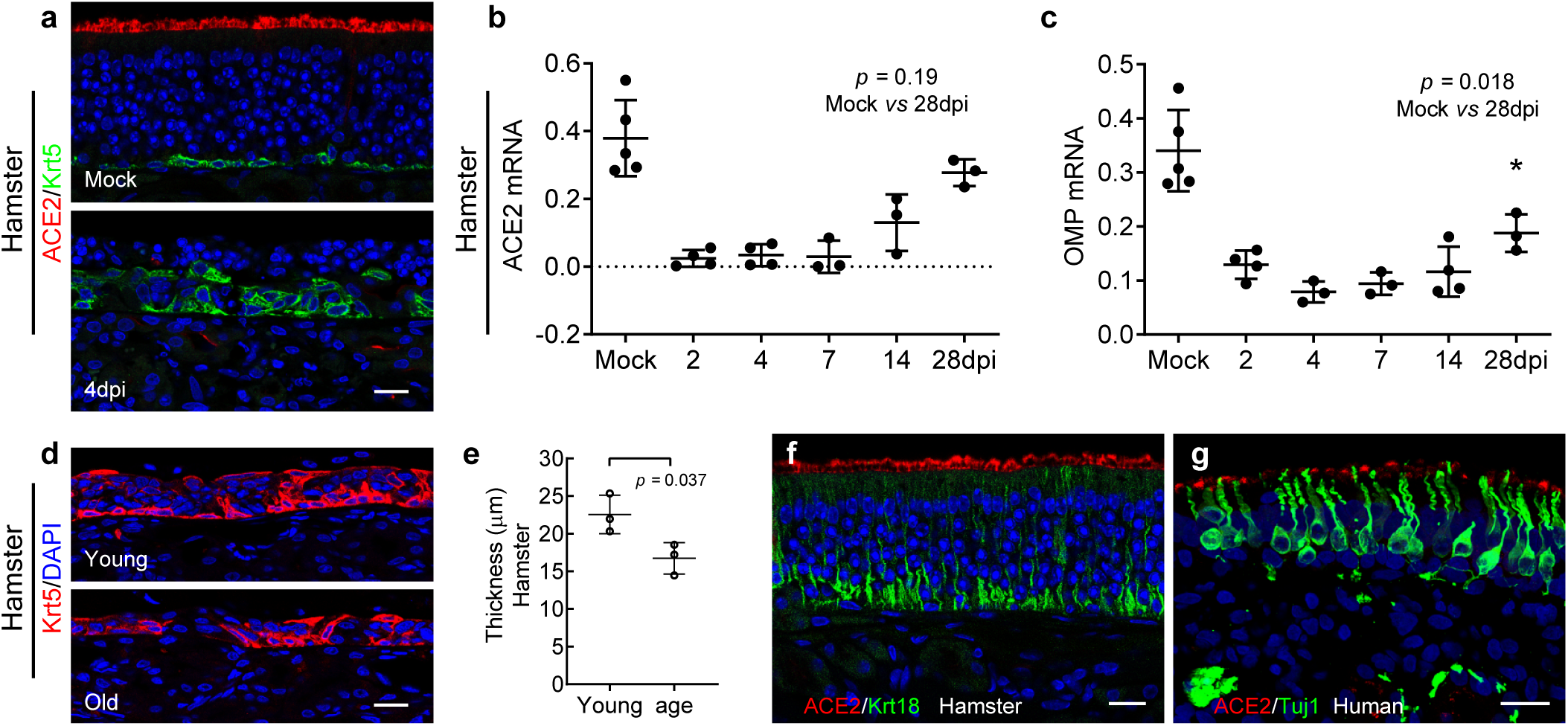
Regeneration of olfactory epithelium and re-expression of ACE2. **a**, Confocal images showing ACE2 (red) and Krt5^+^ horizontal basal cells (green) in olfactory epithelium of mock or SARS-CoV-2 infected hamster at 4 dpi. **b**,**c**, qPCR analysis of ACE2 (**b**) or OMP (**c**) mRNA expression in SARS-CoV-2 infected hamster turbinate lysate at indicated time points. **d**,**e**, Representative images of Krt5^+^ cells in newly regenerated olfactory epithelium (**d**) on 6dpi, and quantification of epithelium thickness (**e**). The thickness of septal olfactory epithelium was measured using Zen lite “line” function. For each section, 8 spots were measured randomly. **f**, Confocal image showing regenerated hamster olfactory epithelium expression of ACE2 at 28dpi. **g**, Representative image shows ACE2 and Tuj1^+^olfactory neurons in an olfactory biopsy from a COVID-19 patient on day 12 post diagnosis. Dots in graph represent independent animal. Data are represented as mean ± S.D. *p* value was calculated by unpaired two-tailed Student’s *t* test. Scale bars, 20 µm.

Because of the limited cross-reactivity of CD45 antibodies with hamster tissue, we took advantage of the mouse adapted SARS-CoV-2 (maSARS) infection model in C57BL/6J wildtype mice^41^. Normally, a low number of CD45^+^ immune cells and Iba1^+^ macrophages/dendritic cells reside in the mouse olfactory mucosa. In maSARS infected group, we observed striking CD45^+^ immune cell infiltration into the lamina propria, crossing the basal cell layer and migrating into the neuroepithelium, suggesting a nasal immune defense in response to viral infection (Fig. 6a). On 6 dpi, approximately 48.3% of CD45^+^ immune cells in olfactory mucosa were Iba1^+^ macrophages/dendritic cells, which is similar to single-cell RNA sequencing data of BALF samples from critical COVID 19 patients^42, 43^.

In hamsters, intranasal inoculation of SARS-CoV-2 induced massive shedding of NP^+^ infected cells into the nasal lumen at 4 dpi (Fig. 6b-d), consistent with our findings in infected human olfactory biopsies. Iba1^+^ macrophages/dendritic cells were widely distributed in the olfactory mucosa and the detached cells in the lumen (Fig. 6b,c). Co-staining analysis showed that the Iba1^+^ macrophages are the major population producing CXCL10 (Extended Data Fig. 7b), a chemokine that has been reported in macrophages from COVID-19 patients’ BALF sample.

Notably, 72% of Iba1^+^ cells were also positive for viral NP antigen at 4dpi (Fig. 6b), indicating uptake of infected cell debris. In addition, some of the apoptotic cells sloughed into the nasal lumen were Iba1^+^/Caspase-3^+^, suggesting the viral clearance by macrophages (Fig. 6c).

Compared to the young hamsters, the number of Iba1^+^ cells in the nasal lumen significantly increased in the older group at 6dpi (Fig. 6d,e). In parallel to the increased macrophages, we observed the number of remaining NP+ cells/debris in the serial sections of older hamster nasal cavities was increased 3.7-fold when compared to young hamsters (Fig. 6d,f), in line with the reported prolonged virus load/delayed viral clearance in older COVID patients^44^. The delayed viral clearance could be a consequence of impaired phagocytic function in aging macrophages, as reported in an influenza infection model^45^.

By analyzing a previously published single cell RNA sequencing (scRNA-seq) dataset^46^ derived from mouse lung CD45^+^ inflammatory cells, we noted significant reduction of phagocytosis related genes^47^ including Clec4n (Dectin2), Fabp5, Fpr2, and Cd9 in old macrophage/dendritic lineages compared to young mice (Fig. 6g). We further verified that the expression of Dectin2 was dominantly located in Iba1^+^ macrophages/dendritic cells in olfactory mucosa of young mice and dramatically declined with age (Fig. 6h). Collectively, our data support that the macrophages are the critical population involved in SARS-CoV-2 defense, and their impaired viral clearance capacity could involve in the prolonged virus retention in the olfactory mucosa of the aged population.

### Regenerated olfactory epithelium regains ACE2 expression

Given the robust reparative capacity of the olfactory mucosa^48^ and the rapid reconstitution post SARS-CoV2 infection^31, 49^, we next systematically examined post-viral stem cell-mediated regeneration using an adult hamster model (2-month old). As a consequence of viral infection, nearly complete loss of neuroepithelium was observed at 4dpi, and ACE2 was not detectable in newly regenerated epithelium (Fig. 7a). Compared to the single layer of Krt5-expressing olfactory stem cells in mock control, SARS-CoV-2 induced widespread epithelial damage and activated robust basal cell proliferation simultaneously (Extended Data Fig. 7c). qPCR analysis revealed that the increased expression of Sox2 (basal cell /sustentacular cell marker), Lgr5 (globose basal cell marker), and Tubb3 (immature neuron marker) was coincident with gradual re-expression of ACE2 as olfactory epithelium regeneration proceeded (Fig. 7b, Extended Data Fig. 7d). The expression of ACE2 and the olfactory sensory neuron marker, OMP, recovered to 78% and 56% of mock on 28 dpi, respectively (Fig. 7b,c). The incomplete recovery of OMP on 28 dpi partially may explain the slow return of olfactory function in human cases with severe damage.

Coincident with epithelial repair, production of CXCL10 in Iba1+ macrophages vanished in both the young and old groups on 6dpi (Extended Data Fig. 7b). Compared to the old group, the newly regenerated olfactory epithelium in young hamsters is significantly thicker at 6dpi (Fig. 7d,e), suggesting age-related delay in regeneration post infection. Furthermore, recovery of ACE2 protein could be detected in hamsters at 28 dpi, and ACE2 expression was also observed in a COVID-19 patient who had lost the sense of smell (Fig. 7f,g).

## Discussion

Understanding the cellular tropism and properties of SARS-CoV-2 infection of the upper airway could provide valuable insights for predicting the pathogenicity of new variants. Consistent with the enrichment of ACE2 in human olfactory sustentacular cells^8^, we herein present greatly enhanced infection efficiency in human and hamster olfactory epithelium, suggesting that this site is potentially critical for initial SARS-CoV-2 infection and replication, especially for the WA1 and Delta strains. The tropism transition from olfactory to respiratory observed in the Omicron variant may explain the low prevalence of anosmia, while the extended duration that Omicron resides in the sinonasal respiratory epithelium may contribute to increased transmission. Our observations, together with the clinical findings of high viral loads in the nasal passages of COVID-19 patients^1, 2^, suggests that the nasal cavity is an important site of SARS-CoV-2 infection, cell damage, and host immune reaction in nasal cavity.

The mechanisms underlying olfactory loss in SARS-CoV-2 infection are difficult to disentangle from a number of pathological processes at multiple anatomic levels^24^.

Quantification of SARS-CoV-2 in nasal and throat swabs reveals a gradual decrease in viral load soon after symptom onset ^2, 50^, suggesting a short pathological process in the nose. Together with these findings, the rapid detachment of infected olfactory epithelium presented here may explain variation in viral loads detected on nasal swabs^2^. The subsequent neuroepithelial structural damage upon viral targeting of supporting sustentacular cells and olfactory neurons plausibly underlies the high incidence of olfactory dysfunction in COVID-19 patients. Importantly, the detached olfactory epithelium likely carries a large amount of virus, and shedding of these infected cells has the potential for aerosolization, exacerbating lung infection, and facilitating transmission between individuals. Other factors include the disrupted nuclear architecture, downregulated olfactory receptor expression^51^ in mild infection, as well as the infection of Bowman’s glands^3^ may also account for the olfactory dysfunction. However, the contribution of the small proportion of olfactory neurons that are become infected based on our observations is likely very limited.

Whether and how SARS-CoV-2 gains access to the brain has been investigated intensively and debated widely^24^. Unlike the obvious infection of the brain in hACE2-expressing mice after SARS-CoV-2 inoculation^52–54^, viral antigen in hamster brain was not detectable^28, 55, 56^ while one study recovered SARS-CoV-2 from brain tissue^55^. A recent study in a hamster model showed limited viral antigen located in nasal OMP^+^ olfactory axons^29^. The presence of SARS-CoV-2 RNA or viral antigen in human postmortem brain tissue reveals that the virus may access the brain even though neuronal infection is rare^25–27^. To avoid the tissue autolysis associated with long postmortem intervals, a bedside endoscopic tissue harvest procedure was developed by Khan et al^3^. In 85 postmortem samples analyzed from COVID-19 cases, even though a uniform sustentacular cell infection was visualized in the olfactory mucosa of a patient within 4 days of diagnosis, no infection in olfactory sensory neurons was identified. It should be noted that the samples in the study by Khan et al were limited to relatively aged (>62 years) patients. Although most children and adolescents are spared from severe COVID-19, it is reported that 22% experience neurologic involvement and 12% develop life-threatening neurologic sequelae^57^.

Abnormal neuroimaging manifestations, including acute disseminated encephalomyelitis-like changes, were also reported in children with COVID-19^58^. Based on infection of young and old hamsters, our observations provide strong evidence that SARS-CoV-2 WA1 targets a subset of mature and immature olfactory neurons, and gains access to the brain through axon transport in an age-dependent manner. The higher proportion of Nrp1^+^ olfactory neuron in the young population may be associated with the increased neuronal infection. It should be noted that a role for other SARS-CoV-2 entry molecules besides Nrp1^59^ for the invasion process cannot be excluded from our data.

The unique targeting of SARS-CoV-2 (WA1 and Delta strains) to a small neuronal population may have impeded discovery to date. As previously mentioned, the absence of evidence for olfactory sensory neuron infection in postmortem samples could be attributed to the older age of the cohort studied^3^. The enhanced olfactory bulb viral transport and subsequent greater level of microglial infiltration in younger hosts may call for a reassessment of neurological impairment in children. Indeed, recent clinical evidence indicates a recurring pattern of disease with SARS-CoV2-related abnormal CNS neuroimaging in infected children without pre-existing conditions^58^. Therefore, the long-term consequences of brain infection require further investigation.

In line with previous observations of aging-related deficits of macrophage phagocytosis in influenza infection models^45^, the delayed SARS-CoV-2 clearance in older hamsters’ olfactory mucosa and in COVID-19 patients may represent a compromised phagocytic function of aged macrophages. The prolonged viral retention may correlate with disease severity in aged COVID-19 patients or with increased risk of transmission. Therefore, the local immune defense in nasal olfactory and respiratory mucosa represents a potential target for early intervention and prevention.

Robust olfactory basal cell activation efficiently regenerates sustentacular cells and restores ACE2 expression. The continued ACE2 expression in the olfactory epithelium may be important, given that anti-SARS-CoV-2 antibodies decay after approximately 6 months from the onset of symptoms, especially in individuals with mild COVID-19 disease^60^. The rapid restoration of ACE2 expression in olfactory epithelium may provide an avenue for re-infection in recovered COVID-19 patients. Taken together, our study identifies the tropism of SARS-CoV-2 WA1 and Delta in olfactory epithelium and the transport of virus to the brain through olfactory neuron axons, especially in younger hosts. In addition, the longer duration of Omicron infection in sinonasal epithelium raises the possibility that early topical intranasal treatment may accelerate viral clearance and reduce transmission.

It should be noted that the observed viral tropism in this study only represents characteristics of infection in the nasal cavity. While our observations demonstrate a high olfactory tropism of WA1 and Delta, the infection is not limited to the olfactory epithelium, and recent RNAseq^61^ and RNAscope or immunohistochemistry^3^ evidence using COVID-19 patient samples suggests the presence of nasal respiratory epithelial infection as well. The extent to which nasal viral load affects lower respiratory infection is not known. In addition, the relatively low amount of virus transported into the olfactory bulb reported here unlikely causes significant neurologic change other than microglial activation and inflammation. Even though the specific cellular tropism in the nasal cavity for each SARS-CoV-2 strain was identified here, it remains to be determined which group of mutations in Spike S protein is associated with altered tropism. Given the predominance of respiratory epithelium by area in the human nasal cavity, the enhanced respiratory infection and the extended viral retention in sinus epithelium may contribute to the increased transmissibility of Omicron, and calls for a reassessment of early local intervention.

---

**Extended Data Fig. 1.**
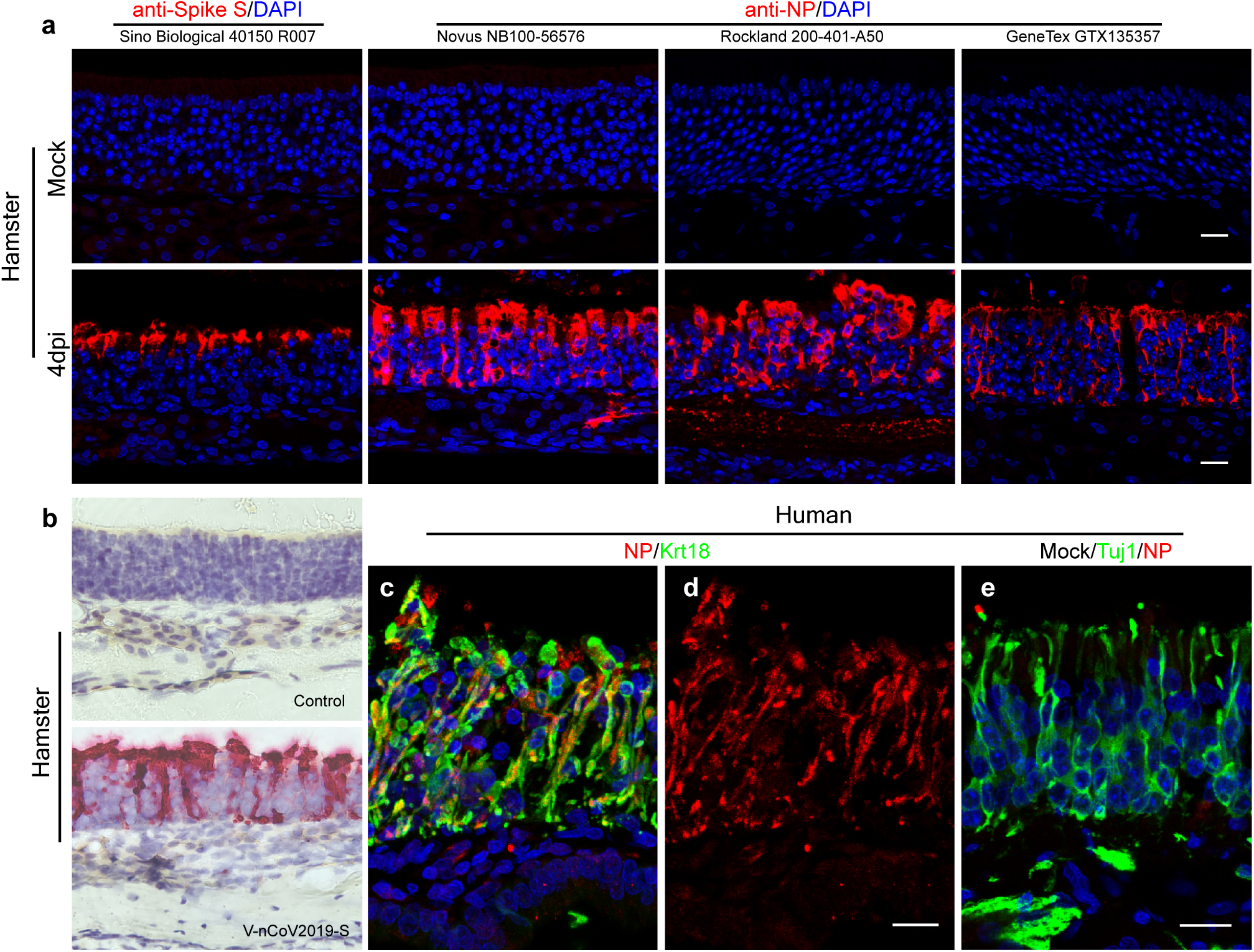
Detection of SARS-Cov-2 in the olfactory neuroepithelium. **a**, SARS-Cov-2 antibody testing. 1 anti-spike S and 3 different anti-NP were verified to be reliable for frozen section immunohistochemistry. Hamster olfactory tissue was examined at 4dpi. All 4 antibodies stained in the same pattern showing intensive viral load mainly located in the apical sustentacular cell layer. No signal could be detected in mock control. Catalog number for each antibody is presented accordingly. **b,** RNAscope analysis showing SARS-Cov-2 viral RNA on 4dpi in hamster olfactory epithelium. **c**,**d**, Co-staining of NP and sustentacular cell marker Krt18. Image was captured from the boxed area in panel (**b**) of Figure 1. **e**, Confocal image of NP and Tuj1 staining in mock control. Scale bars, 20 µm.

**Extended Data Fig. 2.**
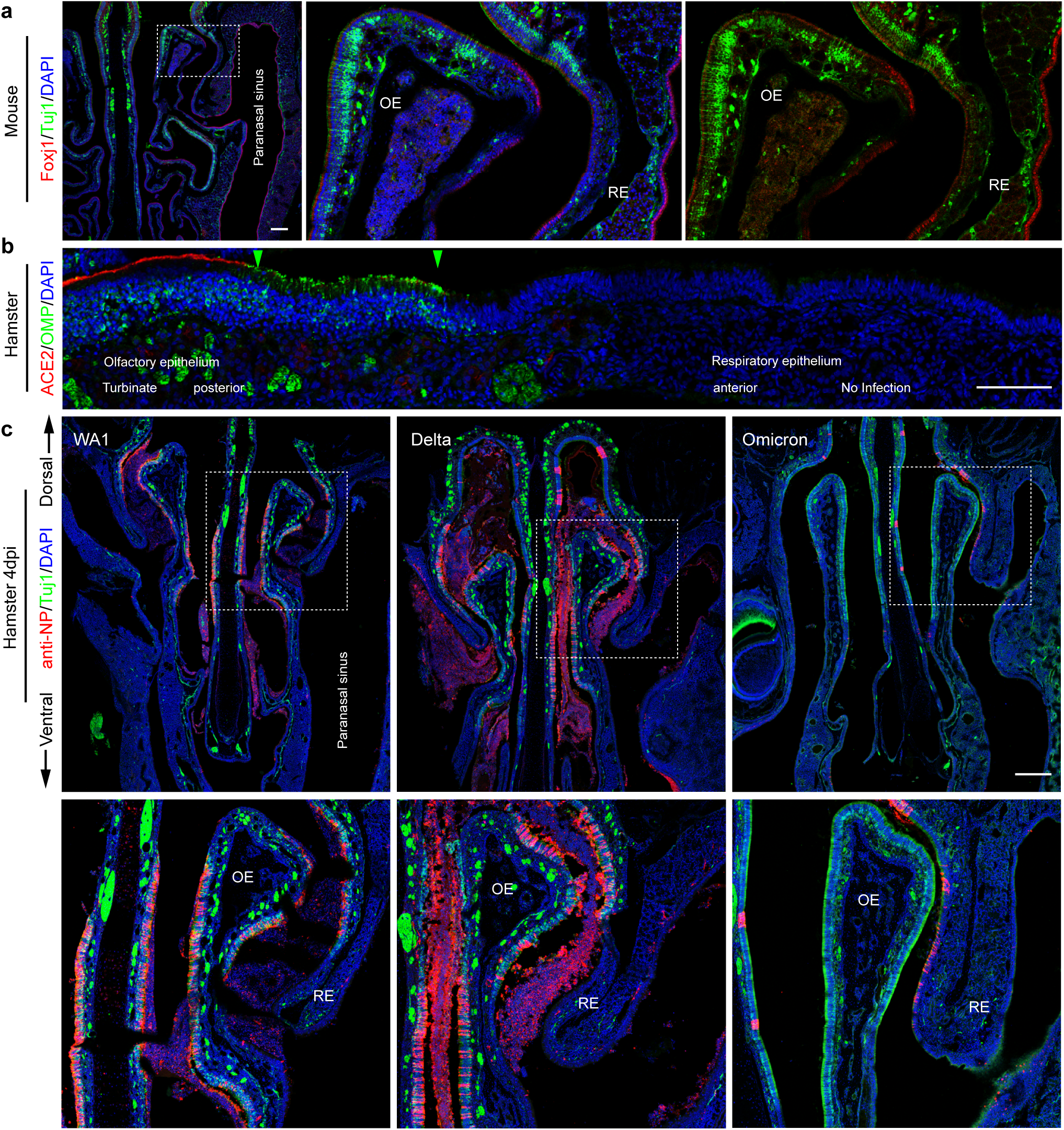
Decreased Omicron variant infection in hamster olfactory epithelium. **a**, Co-staining of neuronal marker Tuj1 and respiratory epithelium marker Foxj1 in mouse nasal cavity. Scale bar, 200 µm. **b**, Representative image shows OMP and rabbit anti ACE2 co-staining in hamster turbinate horizontal section. Intense ACE2 expression is seen in OMP^+^ olfactory epithelium. The green arrows show the respiratory-olfactory transition area with lower ACE2 expression. Scale bar, 100 µm. **c**, Confocal images show the distribution of NP and Tuj1 in a coronal section at L1 of the nasal cavity. Tissues were examined on 4dpi, boxed areas were highlighted at bottom. Note that NP was dramatically declined from Tuj1 negative respiratory epithelium (RE) in hamsters infected with WA1 or Delta. The respiratory infection in Omicron group was markedly increased. Scale bars = 500 µm.

**Extended Data Fig. 3.**
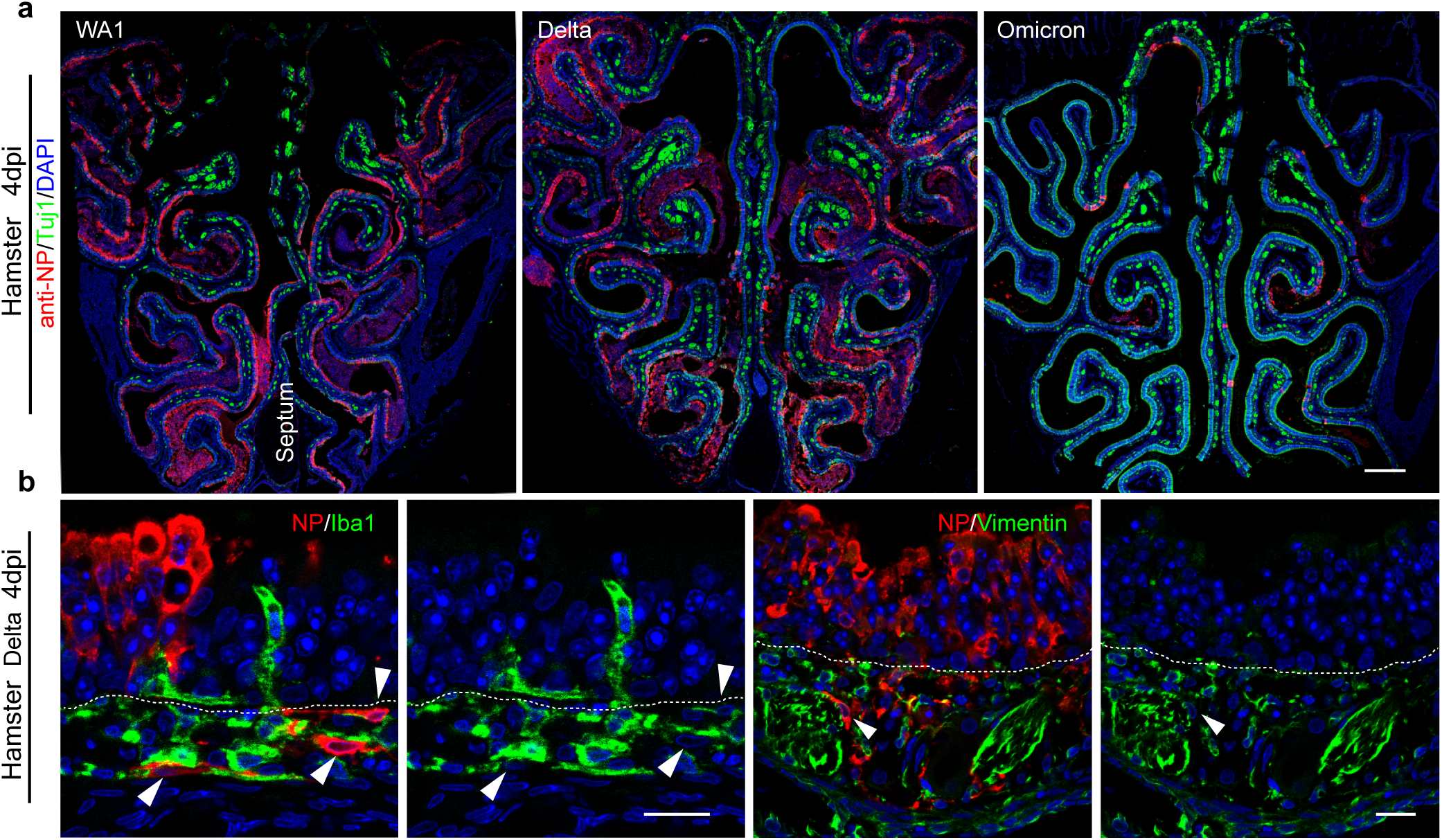
Tropism of SARS-CoV-2 variants in the posterior nasal mucosa. **a**, Confocal images showing the distribution of NP and Tuj1 in posterior nasal cavity sections at 4dpi. Coronal sections at L3 were examined, where the proportion of olfactory epithelium is predominant. The olfactory epithelium infection in the Omicron group was decreased remarkably. Scale bar = 500 µm. **b**, Co-staining of NP and Iba1 (macrophage marker) or NP and Vimentin (mesenchymal cell and olfactory ensheathing cell marker) in Delta infected hamster. Scale bars = 20 µm. The white dotted line in (**b**) indicates the basement membrane.

**Extended Data Fig. 4.**
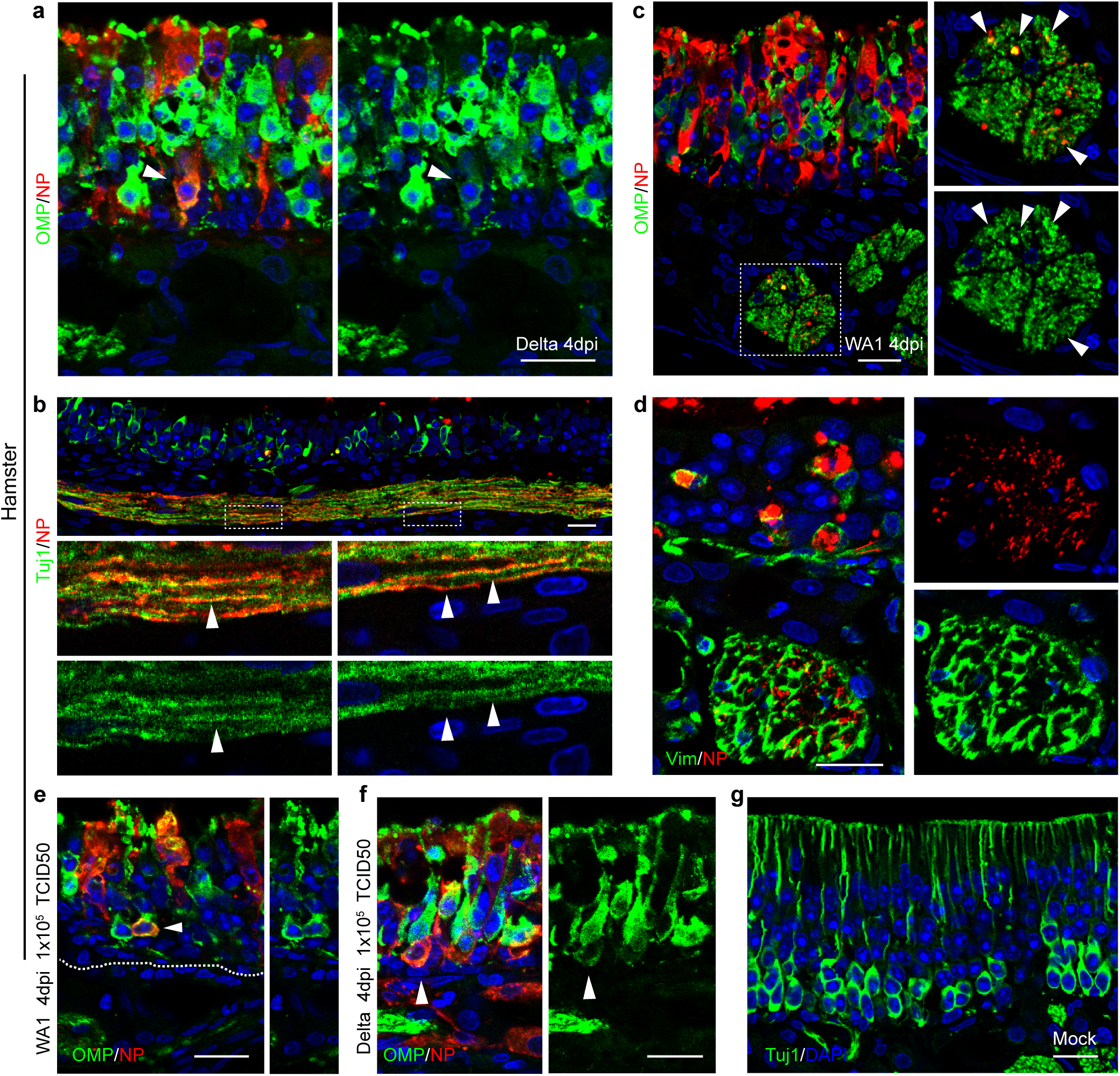
SARS-Cov-2 WA1 and Delta variants infect olfactory sensory neurons. **a**, Co-staining of NP and OMP in olfactory epithelium. 5 weeks-old hamsters were infected with SARS-CoV-2-Delta variant (1 × 10^7^ TCID50) and were examined on 4dpi. Arrowhead indicates an infected neuron. **b**,**c**, Confocal images show co-localization of NP with Tuj1^+^ or OMP^+^ axon. (**b**) shows a larger view of Figure 2f. Boxed areas in (b) were highlighted at bottom. 1m (**b**) or 7-8 weeks-old (**c**) hamsters were infected with WA1. **d**, Representative image shows NP signal does not colocalize with Vimentin in axon bundles. **e**,**f**, Co-staining of NP and OMP in olfactory epithelium. **g**, Confocal image shows Tuj1^+^ immature olfactory neurons in mock group. 7-8 week-old hamsters were infected WA1 (**d, e**) or Delta variant (**f**) at 1 × 10^5^ TCID50 and were examined at 4dpi. Scale bars, 20 µm.

**Extended Data Fig. 5.**
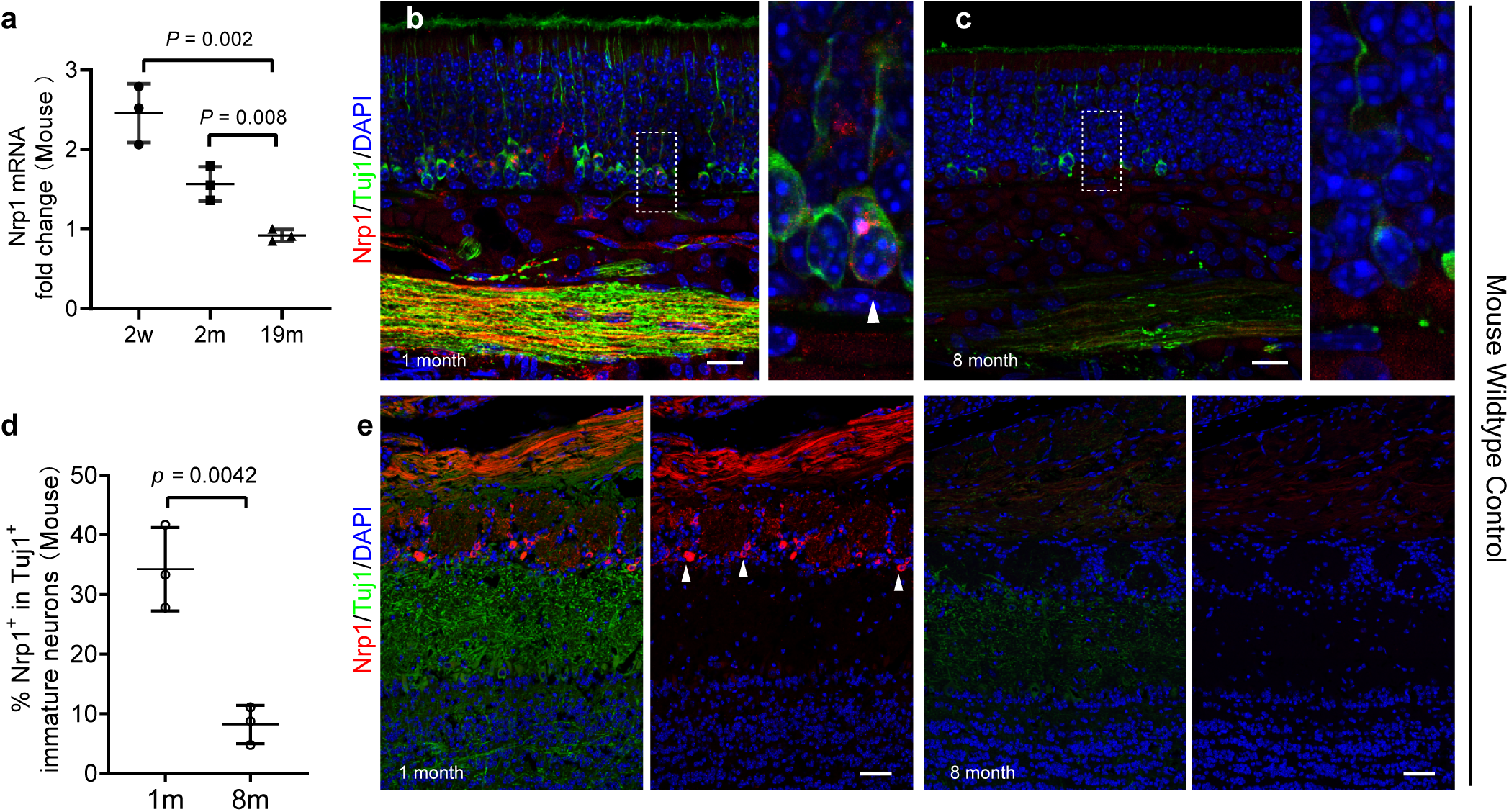
Expression of Nrp1 in mouse olfactory epithelium and bulb. **a**, qPCR analysis of Nrp1 expression in mouse olfactory mucosa at ages 2 weeks, 2 months, and 19 months. Each data point represents an individual mouse (n=3). **b**,**c**, Immunostaining analysis shows the expression of Nrp1 in the Tuj1^+^ immature olfactory neurons and axon bundles. Confocal images were acquired from horizontal section of young (1 month) and old (8 month) mice. Boxed areas were highlighted on right. In young mice, a few mature neurons above the Tuj1^+^ cells also express a low level of Nrp1. **d**, Percentage of Nrp1 expressing cells in Tuj1^+^ immature neurons. Olfactory tissues from wildtype mice were examined at the indicated age groups. **e**, Confocal images show the expression of Nrp1 in young and old mouse olfactory bulb. Data are represented as mean ± S.D. Statistical significance was determined by unpaired two-tailed *t*-test. Arrowheads highlight Nrp1^+^ cells in glomerular layer. Scale bars, 20 µm (**b**,**c**); 50 µm (**e**).

**Extended Data Fig. 6.**
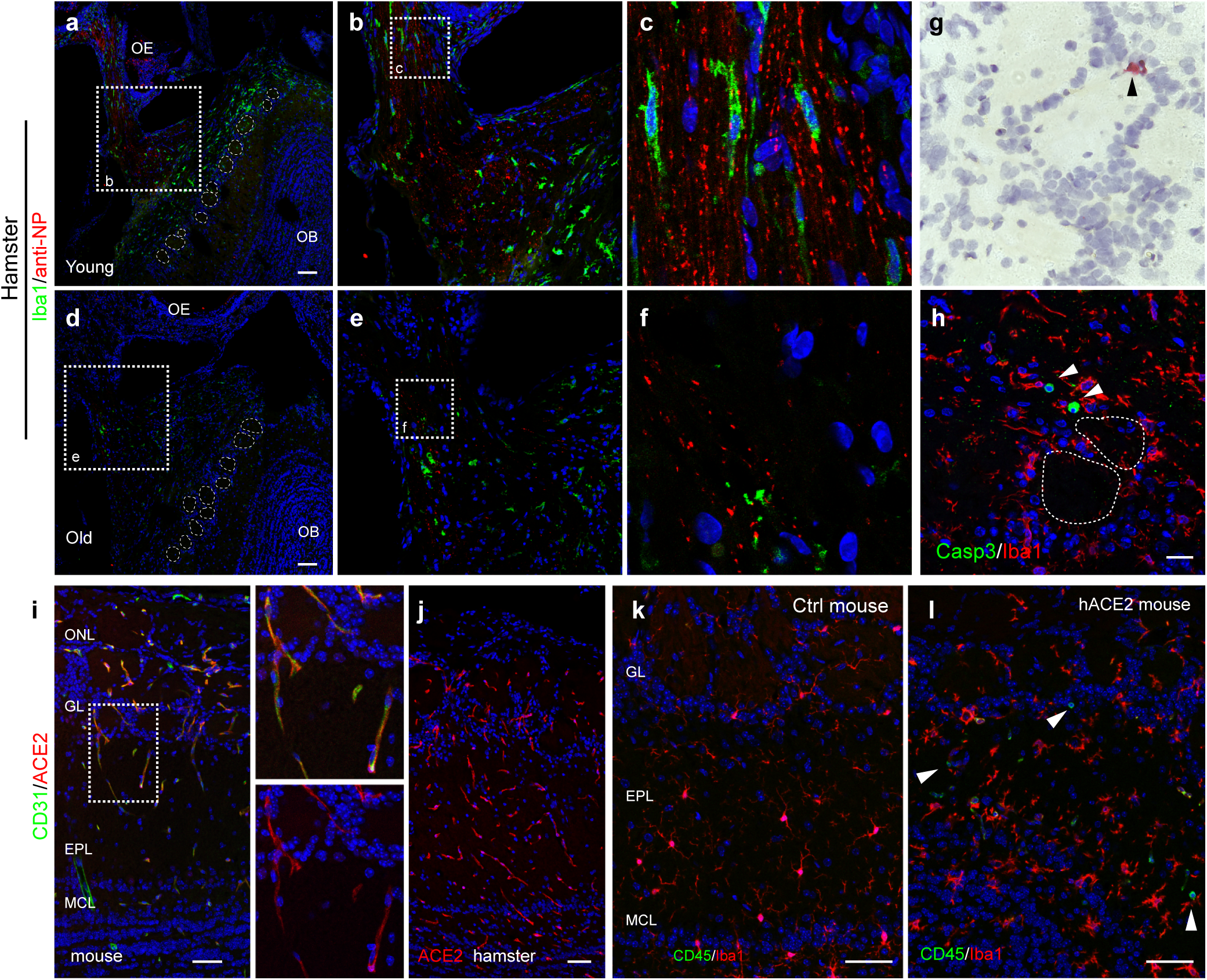
Increased brain transport of SARS-CoV-2 in young hamsters. **a-f**, Confocal image capturing a cross section of olfactory epithelium and olfactory bulb. SARS-CoV-2-infected young or old hamsters were examined at 6 dpi. Boxed areas highlight the infected lateral olfactory axons crossing the cribriform plate and projecting to the olfactory bulb. Images were captured with 4 µm Z-stack and exported by maximum intensity projections. OE, olfactory epithelium; OB, olfactory bulb. **g**, RNAscope analysis shows viral RNA in SARS-Cov-2 infected hamster OB glomeruli at 4dpi. **h,** Co-staining of Caspase-3 and Iba1 in olfactory bulb at 4dpi. **h**,**i**, Confocal image shows co-staining of endothelial cell marker CD31 and ACE2 in mouse (**h**) or hamster (**i,** ACE2 only) olfactory bulb. **j**, Immunostaining of CD45 and microglia marker Iba1 in the olfactory bulb of hACE2 mice at 6 dpi. Arrowheads indicate Iba1 negative immune cells. In the hACE2 strain, human ACE2 overexpression was driven by mouse Krt18 promoter. Scale bars, 100 µm (**a**,**d**); 50 µm (**h**-**j**).

**Extended Data Fig. 7.**
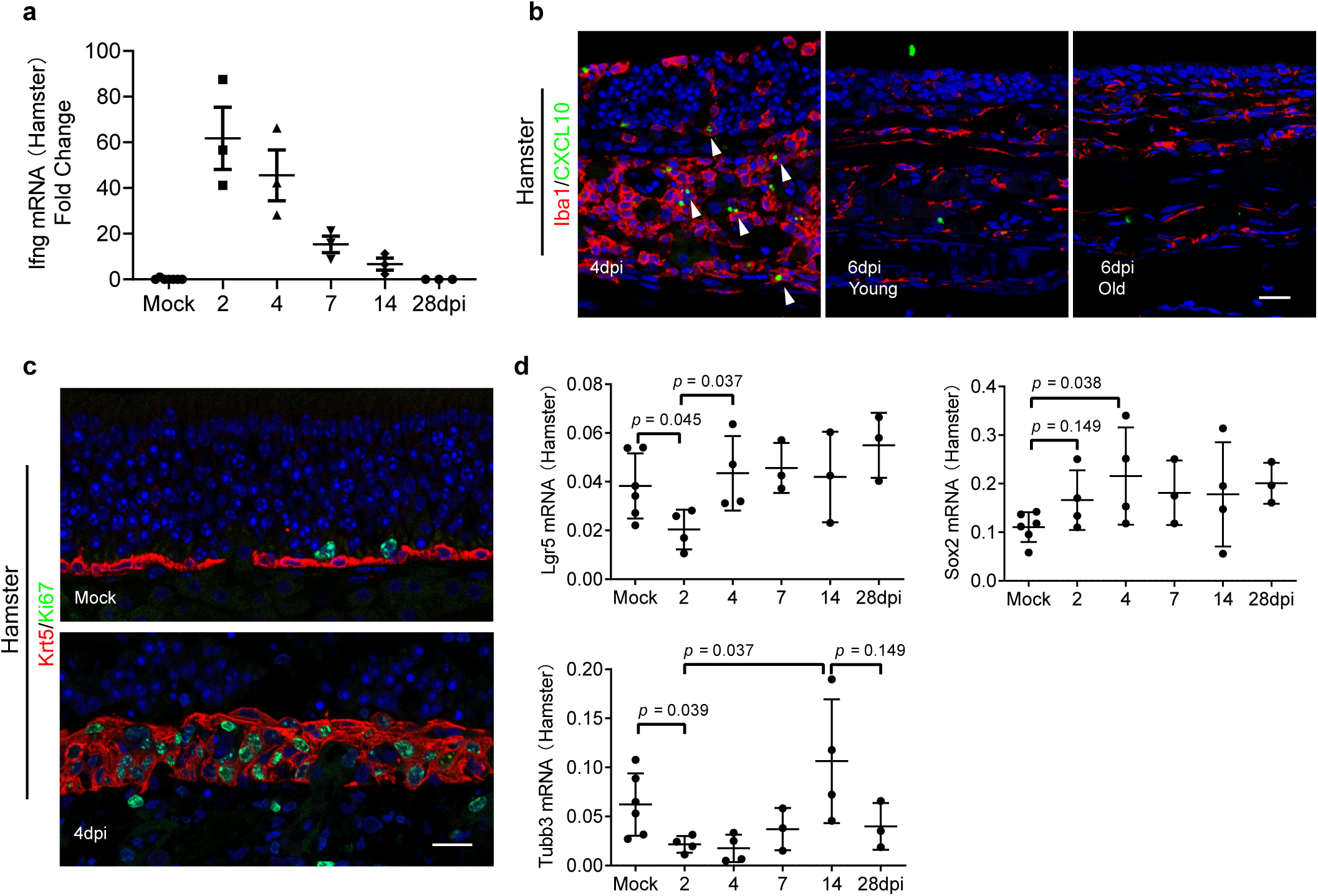
Regeneration of the olfactory epithelium. **a**, qPCR analysis of Ifng expression in turbinate tissues. SARS-CoV-2 infected hamsters were examined at indicated time points. **b**, Dynamic of Iba1^+^ macrophages infiltration and CXCL10 expression in hamster olfactory epithelium. **c**, Representative images show Krt5^+^ basal cells expressing proliferation marker Ki67 on 4dpi in hamster olfactory epithelium. **d**, qPCR analysis of Sox2, Lgr5, and Tubb3 expression in turbinate samples at indicated time points. Data are represented as mean ± S.D. Statistical significance was determined by unpaired two-tailed *t*-test. Each data point represents an individual mouse. Scale bars, 20 µm.

## Methods

### Human nasal explant *in vitro* infection

The research protocol involving human specimens was approved by the Johns Hopkins institutional review board, and all subjects provided signed informed consent. Nasal biopsies included olfactory epithelial and/or respiratory epithelial samples were collected from chronic rhinosinusitis (CRS) patients and control subjects undergoing endonasal surgical approaches for non-CRS disease processes^62^. All patients were tested negative for COVID-19 before surgery. In this study, the majority of biopsies were taken from superior turbinates. The human olfactory mucosa is predominantly distributed on the dorsal aspect of the nasal vault^63^. The superomedial portion of superior turbinate that comprises part of the olfactory cleft contains olfactory epithelium, while the inferior and lateral side is entirely respiratory epithelium. Therefore, the coronal sections of superior turbinate samples in this study include both olfactory and respiratory epithelium, with a much smaller proportion of olfactory relative to respiratory. Notably, over 60% of the superior turbinate biopsies contained solely respiratory epithelium. Other specimens were obtained from the olfactory cleft septal mucosa. More details about the clinical specimens are listed in Supplementary Table 1.

Human biopsies were placed in PneumaCult medium (Stemcell Technologies) and sent for infection immediately. SARS-CoV-2 infection experiments were conducted in a biosafety level 3 facility at the Bloomberg School of Public Health, Johns Hopkins University. After 2 hours incubation with SARS-CoV-2/USA/WA1/2020 (BEI Resources) at 5 × 10^5^ TCID_50_ per mL, the tissues were washed in PBS and transfer into fresh medium at 37°C. Mock controls were maintained in medium without virus. Tissues were fixed at 9 hours post infection in 4% PFA at 4°C for 24 hours. 6 control (2 females and 4 males ranged from 45 to 63 years old) and 27 CRS biopsies (11 females and 16 males ranged from 25 to 76 years old) were used for detailed immunohistochemistry analysis.

Human biopsies for Nrp1 staining were collected from 3 young (20-30 years) and 4 older (68-79years) subjects. Tissues were fixed in 4% PFA at 4°C overnight, and the olfactory neuroepithelium identity was verified by Tuj1 staining.

### Animal *in vivo* infection

Animal infection experiments were carried out in a biosafety level 3 facility at Johns Hopkins Research Animal Resources (RAR) in compliance with the established ethical guidelines.

Animal experimental procedures were approved by the Animal Care and Use Committee at the Johns Hopkins University. Animal infection experiments were conducted using wildtype C57BL/6J mice, Syrian golden hamsters (HsdHan®: AURA, Envigo, Haslett MI), and hACE2 mice (B6.Cg-Tg(K18-ACE2)2Prlmn/J, JAX, Bar Harbor, Maine). In hACE2 strain, the human ACE2 was driven by the mouse Krt18 promoter. 1 × 10^7^ TCID50/ml of SARS-CoV-2/USA/WA1/2020 or 2.4 × 10^7^ TCID50/ml of Delta variant (SARS-CoV-2/USA/MD-HP05660/2021 B.1.617.2) in 100 µL Dulbecco’s modified Eagle medium (DMEM) was intranasally inoculated to hamsters (50 µl per nare). 6.3 × 10^5^ PFU in 20 µL (10 µL per nare) was administered intranasally to hACE2 mice. The mouse-adapted SARS-CoV-2 (courtesy of Michael Schotsaert, Icahn School of Medicine at Mt. Sinai) infection^41^ was performed as 10 µL per nare, 2.5⍰10^8^ PFU. Mock control animals received equivalent volume of DMEM alone.

### Tissue processing

Animals were euthanized in biosafety level 3 facility at indicated time points. After anesthetized with avertin and transcardially perfused with PBS followed by 4% PFA, the skull bone was removed, and the head was postfixed in 4% PFA at 4°C for 3 days. After decalcification in TBD2 solution (6764003, Thermo) overnight and washing in PBS, tissues were equilibrated sequentially in 15% and 30% sucrose, then embedded in Optimum Cutting Temperature (OCT, Tissue-Tek) for sectioning Fixed human biopsies were processed similarly and embedded in OCT without TBD2 treatment. Frozen sections were processed at 12 µm using MICROM HM560 cryostat (Thermo).

### Immunohistochemistry

The immunostaining process was carried out on cryosections after an antigen retrieval step. Briefly, sections were washed in PBS and then blocked in 2% BSA containing 0.2% Triton X-100 at 4 °C overnight, followed by incubation with primary antibodies at 4 °C overnight. The following primary antibodies was used: Rabbit anti-SARS-CoV-2 Nucleoprotein (1:200, Novus, NB100-56576), Rabbit anti-SARS-CoV-2 Nucleoprotein (1:500, GeneTex, GTX135357), Rabbit anti-SARS-CoV Nucleoprotein (1:1000, Rockland, 200-401-A50), Rabbit anti-SARS-CoV-2 Spike S (1:200, Sino Biological, 40150-R007), Goat anti-ACE2 (1:100, R&D, AF933, for human samples), Rabbit anti-ACE2 (1:100, Thermo, MA5-32307, for hamster samples), Goat anti-Neuropilin-1 (1:200, R&D, AF566) Mouse anti-Keratin 18 (1:500, Novus, NB500-306), Goat anti OMP (1:500, Wako, 544-10001), Mouse anti-aSMA (1:100, R&D MAB1420); Chicken anti-Vimentin (Novus NB300-223); Goat anti-Foxj1 (1:200, R&D AF3619); Mouse anti-NeuN (1:1000, BioLegend, 834502); Rat anti-CD45 (1:300, Ebioscience, 14-0451-81), Rat anti-CD31(1:50, BD, 550274), Rabbit anti-Krt5 (1:500, Covance, PRB-160P), Chicken anti-Krt5 (1:500, BioLegend, 905904), Mouse anti Tuj1 (1:300, BioLegend, 801203), Rabbit anti Iba1 (1:500, Wako, 019-19741), Rabbit anti-Cleaved Caspase-3 (1:300, Cell signaling, 9664), Rat anti Dectin2 (1:200, Bio-Rad, MCA2415T), and Goat anti-CXCL10(1:100, R&D, AF-466-NA).

After washing in PBS three times, the tissue sections were incubated with Alexa Fluor conjugated, highly cross-adsorbed, secondary antibodies along with DAPI for nuclear counterstaining. The donkey-derived Alexa Fluor-conjugated secondary antibodies included anti-mouse 488 (A21202, Invitrogen); anti-Rat 488 (A21208, Invitrogen); anti-Rabbit 488 (A21206, Invitrogen); anti-Rabbit 546 (A10040, Invitrogen); anti-Goat 488 (A32814, Invitrogen); anti-Goat 546 (A11056, Invitrogen); anti-Chicken 488 (SAB4600031, Sigma).

### In situ hybridization

To detect SARS-CoV-2 RNA, in situ hybridization was performed on 12 µm-thick sections of 4% PFA-fixed OCT mounted on charged glass slides using the Leica Bond RX automated system (Leica Biosystems, Richmond, IL). Heat-induced epitope retrieval (HIER) was conducted by heating slides to 95°C for 15 minutes in EDTA-based ER2 buffer (Leica Biosystems, Richmond, IL). Slides were treated in protease (Advanced Cell Diagnostics, Newark, CA) for 15 minutes and probes hybridized to RNA for 1 minute. The SARS-CoV-2 probe (#848568, Advanced Cell Diagnostics, Newark, CA) was detected using the Leica RNAScope 2.5 LS Assay-RED kit with a hematoxylin counterstain (#322150, Leica Biosystems, Richmond, IL). An RNApol2 probe served as a host gene control to evaluate RNA quality; a probe for the bacterial dap2 gene served as a negative control ISH probe.

### Confocal Imaging and Quantification

Immunostaining images were obtained using a Zeiss LSM 780 confocal microscope equipped with a 40x, numerical aperture 1.1 water objective. The following laser lines were used DPSS 561nm (detection range 560-612nm) for Alexa Fluor 546; Diode 405nm (detection range 410-480nm) for DAPI; and Argon 488nm (detection range 490-550nm) for Alexa Fluor 488. Images for the same primary antibody across different samples were acquired and exported under the same settings. Before exporting, contrast adjustment was applied as necessary for individual channels using Zen lite (Zeiss) under the “Display” option. Images were cut by Photoshop and assembled by Illustrator.

For quantification, at least 5 images were collected from each specimen using 40x objectives under the tile scan and z stack mode at same depth. Positive cells were identified according to the subcellular staining pattern and were counted manually using “Events” function of Zen lite (Zeiss). Cells in olfactory or respiratory mucosa were quantified per mm of surface epithelium. By measuring the whole length of Tuj1^+^ epithelium, The SARS-CoV2 infected axons were quantified per µm diameter of axon bundle. Microglia in the olfactory bulb or shedding cells in nasal cavity were quantified per mm^2^ tissue.

### RNA isolation, cDNA synthesis and qPCR

Total RNA was isolated from hamster olfactory tissue lysate using a Direct-zol RNA Kits (Zymo). Equal amounts of RNA were transcribed into cDNA by High-Capacity cDNA Reverse Transcription Kit (Applied Biosystems). On-Column DNase I digestion was conducted to remove genomic DNA contamination. Ten nanograms of cDNA was added to a 20-μL PCR reaction using SYBR Green PCR Master Mix or TaqMan Fast Universal PCR Master Mix (Applied Biosystems) on StepOne Plus System (Applied Biosystems). For SYBR Green PCR, post-amplification melting curve analysis was performed to monitor unspecific products. Fold change in mRNA expression was calculated using the comparative cycle method (2^−ΔΔCt^). SYBR Green PCR primer sequences of hamster genes are: ACE2: Forward, TGGTGGGAGATGAAGCGAGA, and Reverse, GAACAGAGCTGCAGGGTCAC; OMP: Forward, CAGAAGCTGCAGTTCGACCG, and Reverse, CAGAAGATTGCGGCAGGGTC; Ifng: Forward, TAATGCACACCACACGTTGC, and Reverse, AAGACGAGGTCCCCTCCATT. GAPDH: Forward, GTGGAGCCAAGAGGGTCATC, and Reverse, GGTTCACACCCATCACAAACAT. Mouse genes are: Nrp1: Forward, CAGTGGCACAGGTGATGACT, and Reverse, ACCGTATGTCGGGAACTCTGAT; ACE2: Forward, CCATTGGTCTTCTGCCATCCG, and Reverse, CCAACGATCTCCCGCTTCATC; GAPDH: Forward, TCAATGAAGGGGTCGTTGAT, and Reverse, CGTCCCGTAGACAAAATGGT.

### Single cell RNA-seq analysis

Sc-RNA-seq dataset was retrieved from published study (GSE155006) by Mogilenko, et al^46^. This dataset was generated from sorted lung CD45^+^ immune cells from 3 or 17-month-old mice. The Seurat R package was used for subsequent analysis. Quantity control was conducted according to the standard pre-processing workflow. Cells in young and old datasets express 500-2500 genes, mitochondrial genes less than 5% were selected and normalized using a scaling factor 10,000. The highly variable genes in each dataset were selected using the *FindVariableFeatures*function, and combined (10,228 cells in total) for Seurat integration procedure and linear dimensionality reduction. The top 2000 most variable genes per dataset were used for downstream principal component analysis and clustered using the *FindClusters* function. The datasets include 16 clusters were then projected as UMAP plots. According to the expression levels of canonical marker genes, we matched the clusters to known immune cell types. We applied *FindMarkers* function to identify differentially expressed genes in macrophages/dendritic cell lineage between young and old conditions. Average Log_2_ fold changes of gene expression and the percentage of cells expressing certain genes in each condition were calculated.

### Statistical analyses

Data are expressed as mean ± SD. as indicated. Data analyses were carried out using GraphPad Prism. For experiments with two groups, *P* values were calculated using the unpaired two-tailed Student’s t-test. Differences were considered significant when *P* < 0.05.

### Reporting Summary

Further information on research design is available in the Nature Research Reporting Summary linked to this article.

### Data and materials availability

Data, code, and materials will be made available upon request.

## Acknowledgements

We are grateful to patients consenting to donate specimens for research. We thank Andrew Johanson and Riley Richardson for RNAscope analysis. These studies were supported through the generosity of the collective community of donors to the Johns Hopkins University School of Medicine for COVID research. This work was funded by NIH Grants R01 AI132590, R01 DC016106 (A.P.L) and NIAID HHS N2772201400007C (AP).

## Author contributions

M.C., A.P.L., and A.P. designed research; M.C. and W.S. performed experiments; M.C. performed the bioinformatic analysis of the single cell RNA-seq dataset; A.P., R.Z. performed *in vitro* infection experiments; N.R.R., A.P.L., M.R., H.K., and Z.L. collected biopsies; J. V., S. B., and J. M. performed animal infection experiments; M.C., A.P., J. M., K. W. and A.P.L. analyzed data; M.C., A.P., and A.P.L. wrote the paper.

## Competing interests

The authors declare no competing interests.

## Notes

https://livejohnshopkins-my.sharepoint.com/:f:/g/personal/mchen85_jh_edu/Ek_raEjP8DNCuqZ7WPUpM0UBT44K6D1vP-mN22qOIuFzvA?e=QEoF8O

### Competing Interest Statement

The authors have declared no competing interest.

