## Supplementary material for "Evolution of nasal and olfactory infection characteristics of SARS-CoV-2 variants": Supp Table 1

Supplementary Table 1. Characters of human olfactory epithelium biopsies

| No | Gender | Age | Condition | Biopsy Location | Figure | OE/RE | Infection or Nrp1 staining |
|---|---|---|---|---|---|---|---|
| 072220 | F | 33 | CRS | ST | Fig 1a | OE+RE | Infection |
| 072820 | M | 25 | CRS | ST | Fig 1g | OE+RE | Infection |
| 081220 | F | 55 | CRS | ST | Fig 1g | OE+RE | Infection |
| 090920 | M | 68 | CRS | ST | Fig 1g | OE+RE | Infection |
| 101920 | F | 72 | CRS | ST | Fig 1b | OE+RE | Infection |
| 091620 | F | 30 | CRS | ST | Fig 1g | OE+RE | Infection |
| 092120 | M | 54 | Control | ST | Fig 1g | OE+RE | Infection |
| 092120 | M | 54 | Control | Olfactory cleft | Fig 1f | OE | Infection |
| 091420 | F | 36 | Control | ST | Fig 1d | OE+RE | Mock |
| 100520 | M | 41 | Control | ST | NS | OE+RE | Mock |
| 081020 | M | 50 | Control | ST | NS | OE+RE | Mock |
| 061820 | F | 59 | CRS | ST | Fig 1g | RE | Infection |
| 062620 | F | 73 | Control | ST | Fig 1g | RE | Infection |
| 070820 | M | 67 | CRS | ST | Fig 1g | RE | Infection |
| 071620 | M | 48 | CRS | ST | Fig 1g | RE | Infection |
| 071720 | F | 71 | CRS | ST | Fig 1g | RE | Infection |
| 072120 | F | 58 | Control | ST | Fig 1g | RE | Infection |
| 072220 | M | 57 | CRS | ST | Fig 1g | RE | Infection |
| 073120 | F | 36 | Control | ST | Fig 1g | RE | Infection |
| 080520 | M | 58 | CRS | ST | Fig 1g | RE | Infection |
| 082620a | F | 57 | CRS | ST | Fig 1e,g | RE | Infection |
| 082620b | F | 62 | CRS | ST | Fig 1g | RE | Infection |
| 090220a | F | 56 | Control | ST+Septum | Fig 1g | RE | Infection |
| 090220b | M | 26 | Control | ST | Fig 1g | RE | Infection |
| 091420 | M | 72 | CRS | ST Left+Right | Fig 1g | RE | Infection |
| 091520a | F | 61 | CRS | ST | Fig 1g | RE | Infection |
| 091520b | M | 46 | CRS | ST | Fig 1g | RE | Infection |
| 091620 | M | 34 | CRS | ST | Fig 1g | RE | Infection |
| 093020 | F | 65 | CRS | ST | Fig 1g | RE | Infection |
| 102620 | M | 48 | CRS | ST | Fig 1g | RE | Infection |
| 101220 | F | 68 | Control | ST | Fig 1g | RE | Infection |
| 110220 | M | 63 | CRS | ST | Fig 1g | RE | Infection |
| 110420 | M | 77 | CRS | ST | Fig 1g | RE | Infection |
| 111020 | F | 41 | CRS | ST | Fig 1g | RE | Infection |
| 111320 | M | 65 | CRS | ST+Sinus | Fig 1g | RE | Infection |
| 040721 | F | 27 | Control | ST | Fig 1j | OE+RE | Nrp1 |
| 021721 | F | 30 | CRS | ST | Fig 1j,k | OE+RE | Nrp1 |
| 050721 | M | 20 | CRS | ST | Fig 1j | OE+RE | Nrp1 |
| 022221a | M | 68 | CRS | ST | Fig 1j | OE+RE | Nrp1 |
| 022221b | F | 69 | Control | ST | Fig 1j | OE+RE | Nrp1 |
| 090820 | F | 72 | CRS | ST | Fig 1j,k | OE+RE | Nrp1 |
| 040521 | M | 79 | Control | ST | Fig 1j | OE+RE | Nrp1 |

OE: olfactory epithelium; RE, respiratory epithelium; ST, superior turbinate; NS, image not shown.
